# A microbiota-derived metabolite instructs peripheral efferocytosis

**DOI:** 10.1101/2022.08.17.504322

**Authors:** Pedro H. V. Saavedra, Alissa Trzeciak, Zhaoquan Wang, Waleska Saitz Rojas, Giulia Zago, Melissa D. Docampo, Jacob G. Verter, Marcel R. M. van den Brink, Christopher D. Lucas, Christopher J. Anderson, Alexander Y. Rudensky, Justin S. A. Perry

**Affiliations:** Immunology Program, Sloan Kettering Institute, Memorial Sloan Kettering Cancer Center, New York, NY, USA; Immunology and Microbial Pathogenesis Program, Weill Cornell Medical College, Cornell University, New York, NY, USA; Adult Bone Marrow Transplantation Service, Department of Medicine, Memorial Sloan Kettering Cancer Center, New York, NY, USA; Department of Medicine, Weill Cornell Medical College, Cornell University, New York, NY, USA; University of Edinburgh Centre for Inflammation Research, Queen’s Medical Research Institute, Edinburgh BioQuarter, UK; Institute for Regeneration and Repair, Edinburgh BioQuarter, UK; VIB-UGent Center for Inflammation Research, Ghent, Belgium; Department of Biomedical Molecular Biology, Ghent University, Ghent, Belgium; Howard Hughes Medical Institute and Ludwig Center, Memorial Sloan Kettering Cancer Center, New York, NY, USA; Louis V. Gerstner Jr. Graduate School of Biomedical Sciences, Memorial Sloan Kettering Cancer Center, New York, NY, USA

## Abstract

The phagocytic clearance of dying cells, termed efferocytosis, is essential for both tissue homeostasis and tissue health during cell death-inducing treatments. Failure to efficiently clear dying cells augments the risk of pathological inflammation and has been linked to a myriad of autoimmune and inflammatory diseases. Although past studies have elucidated local molecular signals that regulate efferocytosis in a tissue, whether signals arising distally also regulate efferocytosis remains elusive. Interestingly, clinical evidence suggests that prolonged use of antibiotics is associated with an increased risk of autoimmune or inflammatory disease development. We therefore hypothesized that intestinal microbes produce molecular signals that regulate efferocytotic ability in peripheral tissue phagocytes. Here, we find that macrophages, the body’s professional phagocyte, display impaired efferocytosis in peripheral tissues in both antibiotic-treated and germ-free mice *in vivo*, which could be rescued by fecal microbiota transplantation. Mechanistically, the microbiota-derived short-chain fatty acid butyrate directly boosted efferocytosis efficiency and capacity in mouse and human macrophages, with both intestinal and local delivery of butyrate capable of rescuing antibiotic-induced peripheral efferocytosis defects. Bulk mRNA sequencing of primary macrophages treated with butyrate *in vitro* and single cell mRNA sequencing of macrophages isolated from antibiotic-treated and butyrate-rescued mice revealed specific regulation of phagocytosis-associated transcriptional programs, in particular the induction of programs involved in or supportive of efferocytosis. Surprisingly, the effect of butyrate on efferocytosis was not mediated through G protein-coupled receptor signaling, but instead acted by inhibition of histone deacetylase 3. Strikingly, peripheral efferocytosis was impaired well-beyond withdrawal of antibiotics and, importantly, antibiotic-treated mice exhibited a poorer response to a sterile efferocytosis-dependent inflammation model. Collectively, our results demonstrate that a process essential for tissue homeostasis, efferocytosis, relies on distal molecular signals, and suggest that a defect in peripheral efferocytosis may contribute to the clinically-observed link between broad-spectrum antibiotics use and inflammatory disease.

## Introduction

Over 1% of our cellular biomass, approximately 3×10^11^ cells, is turned over every day (*1*). The majority of this cell turnover is performed by phagocytes embedded in tissues, such as tissue-resident macrophages, through a process termed efferocytosis (*2*–*5*). Beyond homeostatic efferocytosis, clearance of dying cells also facilitates resolution of inflammation and return to homeostasis when a tissue is damaged via sterile injury. Indeed, failure to efficiently clear dead cells via efferocytosis is associated with the onset and progression of numerous autoimmune and inflammatory diseases (*2*, *4*, *6*–*10*). Past studies have focused on both the signals produced within a tissue such as growth factors or metabolites, and local cell populations that regulate efferocytosis (*11*–*13*). However, how molecular signals generated outside of a given tissue (‘distal signals’) regulate a phagocyte’s efferocytosis potential remain unexplored (*14*). This notion, that distal signals regulate efferocytosis, is supported by the finding that efferocytosis, at least in some contexts, is facilitated by Protein S (*15*). Protein S is found in serum and can be generated both by cells locally and in the liver (*16*). Protein S, which acts by binding to phosphatidylserine on an apoptotic cell, constitutes a putative example of a distal signal that directly facilitates recognition and internalization of an apoptotic cell (*16*). On the other hand, whether there exist distal signals that act directly on a phagocyte to regulate efferocytosis remains unknown.

Mammals co-evolved with a microbiome in a symbiotic relationship that ensures homeostasis of both host and symbiote. The importance of this relationship is illustrated by the numerous examples of dysregulated microbiome (dysbiosis) negatively affecting host physiological processes (*17*–*19*) and is exemplified by the production of short-chain fatty acids (SCFAs). Specifically, intestinal bacteria capture components from the host diet for their own metabolic needs, but in the process produce SCFAs. SFCAs, in turn, have been shown to regulate the homeostatic functions of both immune and non-immune cells in the intestines (*20*, *21*). SCFAs, in particular butyrate, are found at high levels in the serum (~80μmol/l) and are strong candidates for distal signals given past evidence suggesting their role in non-intestinal tissues (*22*, *23*). Interestingly, numerous clinical and epidemiological studies suggest that the use of antibiotics increases the risk of developing autoimmune or inflammatory disease (*24*–*32*). Given the importance of the intestinal microbiome to host homeostasis, its known role in immune cell regulation, and the direct link between antibiotics-induced dysbiosis and onset of diseases related to impaired efferocytosis, we reasoned that the molecular signals produced by the microbiome regulate efferocytosis by peripheral tissue phagocytes.

## Results

### The intestinal microbiome supports peripheral efferocytosis

To investigate whether healthy intestinal microbiota informs the ability of peripheral tissue-resident macrophages (TRMs) to perform efferocytosis, we focused on peritoneal macrophages (pMacs), a highly phagocytic, long-lived resident TRM population with direct access to circulating substrates (*33*, *34*). We treated mice with an oral course of broad-spectrum antibiotics prior to peritoneal injection of apoptotic cells *in vivo* (**Fig. 1A**). Analysis of peritoneal exudates revealed that resident CD11b^+^ F4/80^+^ macrophages (Fig. S1A), the phagocyte primarily responsible for efferocytosis in the peritoneum (*35*), exhibit dramatically decreased ability to internalize and digest apoptotic cells in mice treated with a 7d course of antibiotics (**Fig. 1A**), corresponding to the timepoint at which the majority of intestinal bacteria are removed (*36*). Antibiotics have been shown to directly affect phagocytosis of different targets including apoptotic cells in culture, albeit generally increasing, not decreasing, phagocytosis capacity (*37*–*39*). As an alternative approach to antibiotics, we performed peritoneal efferocytosis assays with germ-free (GF) mice. Compared to specific-pathogen-free (SPF) mice, CD11b^+^ F4/80^+^ pMacs exhibited significantly decreased efferocytosis capacity in GF mice (**Fig. 1B**), without significant changes in pMac numbers (Fig. S1B to C). Thus, mice lacking intestinal microbes exhibit defective efferocytosis in a peripheral tissue.

**Fig. 1.**
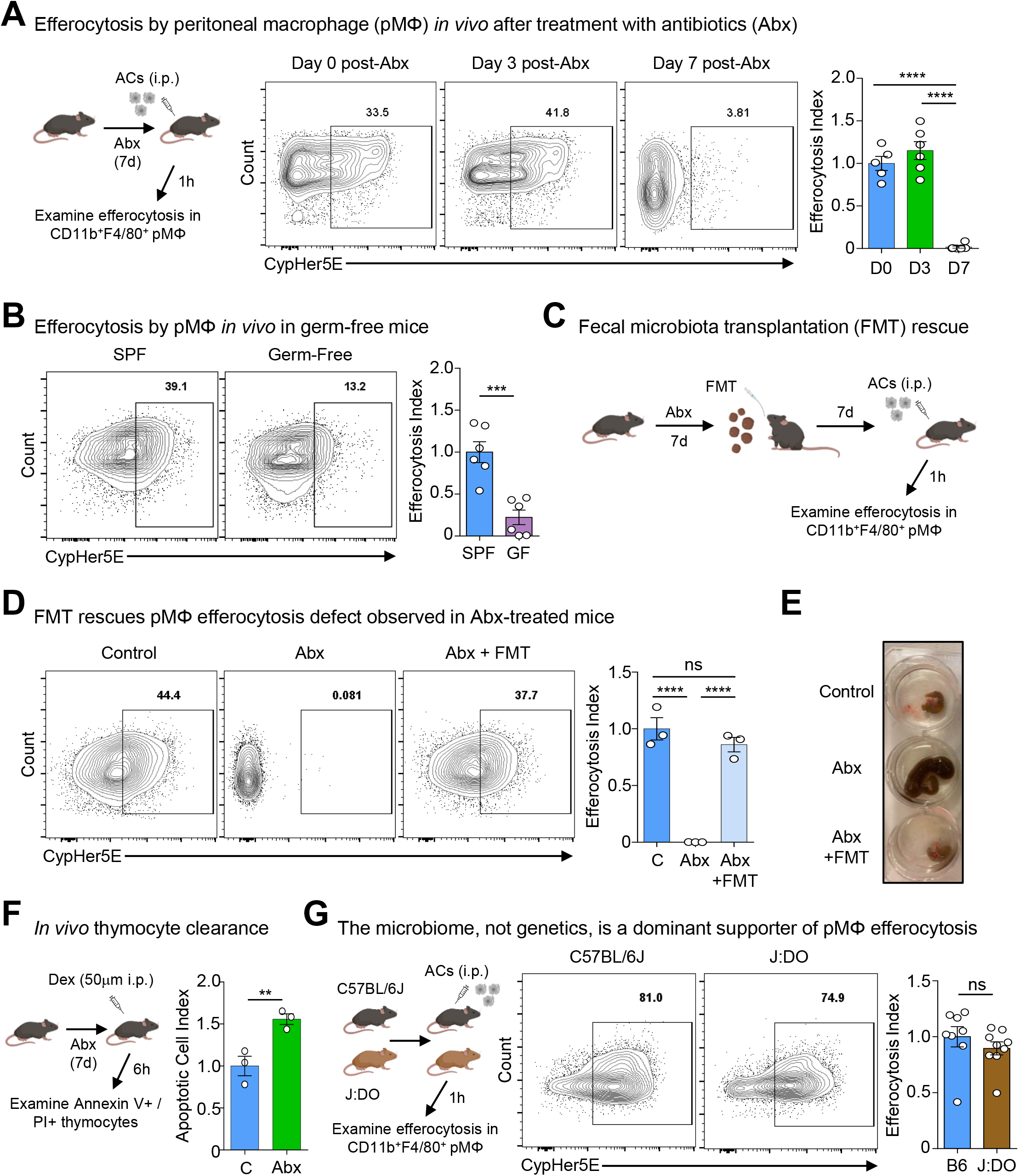
The intestinal microbiome supports peripheral efferocytosis. (A) <u>Peritoneal macrophages exhibit decreased efferocytosis capacity in antibiotic-treated mice *in vivo*</u>. (left) Schematic of experimental design. Mice were treated with antibiotics in drinking water for 0-7d prior to intraperitoneal (i.p.) injection of CypHer5E-labeled apoptotic cells (ACs). After 1h, peritoneal CD11b^+^F4/80^+^ macrophages were analyzed for efferocytosis. Shown are representative flow cytometry (center) and summary (right) plots of the rate of efferocytosis by CD11b^+^F4/80^+^ macrophages in untreated mice (D0, n = 5), in mice after 3d of antibiotics (D3, n = 6), and in mice after 7d antibiotics (D7, n = 6). \*\*\*\**p* < .0001. Data are from two independent experiments with between two to three mice per experiment. (B) <u>Peritoneal macrophages exhibit decreased efferocytosis capacity in germ-free mice *in vivo*</u>. *In vivo* efferocytosis experiments were performed similar to (A), but in specific pathogen-free (SPF) and germ-free (GF) mice. Shown are representative flow cytometry (left) and summary (right) plots of the rate of efferocytosis by CD11b^+^F4/80^+^ macrophages in SPF (n = 6) and GF (n = 6) mice. \*\*\**P* < .001. Data are from two independent experiments with three mice per experiment. (C-E) <u>Microbiome reconstitution via fecal microbiota transplantation (FMT) rescues efferocytosis capacity by peritoneal macrophages in antibiotic-treated mice *in vivo*</u>. (C) Schematic of experimental design. Mice were treated with antibiotics in drinking water for 7d. Some antibiotic-treated mice were supplemented via fecal microbiota transplantation (FMT) of homogenized stool from SPF mice. After 7d, mice were injected i.p. with CypHer5E-labeled ACs. After 1h, peritoneal CD11b^+^F4/80^+^ macrophages were analyzed for efferocytosis. (D) Shown are representative flow cytometry (left) and summary (right) plots of the rate of efferocytosis by CD11b^+^F4/80^+^ macrophages in control mice (n = 3), in antibiotic-treated mice (ABX, n = 3), and in antibiotic-treated mice supplemented via FMT (ABX+FMT, n = 3). (E) Shown are representative images of control mice, antibiotic-treated mice, and antibiotic-treated mice supplemented via FMT. \*\*\*\**p* < .0001. ns = not significant. Data are from two independent experiments with between one to two mice per experiment. (F) <u>Thymic efferocytosis is impaired in antibiotic-treated mice *in vivo*</u>. Schematic of experimental design (left) and summary plot (right) of AnnexinV^+^PI^+^ from isolated thymocytes 6h post-dexamethasone (Dex) injection in vehicle (n = 3) or antibiotic-treated mice (n = 3). \*\**p* < .01. Data are from three experimental replicates. (G) <u>Peritoneal macrophage efferocytosis capacity is informed primarily by the microbiome, not genetics.</u> (left) Schematic of experimental design. Similar to (A), C57BL6/J or J:DO mice were injected i.p. with CypHer5E-labeled ACs. After 1h, peritoneal CD11b^+^F4/80^+^ macrophages were analyzed for efferocytosis. Shown are representative flow cytometry (middle) and summary (right) plots of the rate of efferocytosis by CD11b^+^F4/80^+^ macrophages in C57BL6/J (B6, n = 8) and J:DO (n = 9) mice. ns = not significant. Data are from two independent experiments with between four to five mice per experiment.

Both antibiotic-treated and GF mice exhibit sequelae not necessarily caused by absence of intestinal microbes (*40*–*45*). To directly test the requirement of intestinal microbes on efferocytosis in the peritoneum, we reconstituted the microbiomes of antibiotic-treated mice via fecal microbiome transplantation (FMT) (**Fig. 1C**). Importantly, antibiotic-treated mice reconstituted via FMT exhibited restored efferocytosis capacity (**Fig. 1D**) and concomitant reversal of the enlarged caecum observed in antibiotic-treated mice (**Fig. 1E**), indicating efficient intestine recolonization. Our finding that efferocytosis by peritoneal macrophages was restored in FMT recipients suggested that factors originating from intestinal microbes continually educate resident pMacs instead of imprinting multipotent myeloid progenitors or hematopoietic stem cells, as has been observed in different contexts (*46*–*48*). In support of the former, we found that macrophages differentiated from stem/progenitor cells isolated from GF and SPF mice exhibited equivalent efferocytosis capacity (Fig. S1D). Lastly, we sought to determine if the defect in efferocytosis observed in the peritoneum extends to other peripheral tissues. To this end, we used the dexamethasone-induced thymocyte cell death model to test efferocytosis in the thymus. In this model, dexamethasone induces a wave of thymocyte apoptosis which is subsequently cleared by thymic macrophages, with accumulation of uncleared apoptotic cells indicative of deficient efferocytosis (*35*). Similar to our findings in the peritoneum, we found that antibiotic-treated mice exhibit significantly decreased efferocytosis capacity (as indicated by a significant increase in uncleared apoptotic cells; **Fig. 1F** and Fig. S1E).

We next queried whether microbiome-derived signals were a dominant factor informing peripheral efferocytosis capacity by TRMs. To test this, we took advantage of the Jackson Laboratory-maintained Diversity Outbred (J:DO) strain, which share similar microbiomes with JAX-obtained C57BL/6J mice but exhibit highly heterogeneous genomes across individuals (*49*). We housed J:DO and age- and sex-matched C57BL/6J mice in our animal facility for ~4 weeks prior to performing peritoneal efferocytosis assays to further normalize microbial composition. Fascinatingly, we observed similar efferocytosis capacity between J:DO and C57BL/6J mice (**Fig. 1F**). We did however observe that J:DO mice displayed an increase in the fraction of CD11b^+^ F4/80^+^ macrophages (Fig. S1F). Because our *in vivo* efferocytosis assay is performed with saturating numbers of apoptotic cells, an elevation in the frequency of macrophages is unlikely to confound the interpretation of the current experiment. However, in tissues or contexts where apoptotic cell numbers are limiting or sparsely distributed, an abundance of mature TRMs may provide an added layer of protection against a breakdown in homeostasis. Taken together, our findings suggest that intestinal microbe-derived factors are a dominant signal informing efferocytosis by tissue-resident macrophages in the periphery.

### Butyrate boosts efferocytosis via induction of efferocytotic transcriptional programs

Given our finding that dysbiosis impairs efferocytosis by peritoneal macrophages (pMacs) and that microbiota-derived metabolites such as short-chain fatty acids (SCFAs) regulate immune cell function locally (*50*) and possibly peripherally (*51*, *52*), we sought to determine if SCFAs affect peripheral macrophage efferocytosis. To this end, we first cultured stem/ progenitor cells with one of the SCFAs butyrate, acetate, or propionate during macrophage differentiation (Fig. S2A). Treatment with butyrate, but not acetate or propionate, markedly increased efferocytosis efficiency (**Fig. 2A**) and capacity (**Fig. 2B**) compared to vehicle-treated macrophages. Importantly, mature macrophages treated with butyrate also significantly upregulated efferocytosis efficiency and capacity compared to vehicle-treated macrophages (**Fig. 2C**), consistent with our *in vivo* data suggesting that intestinal microbiota-derived factors act directly on mature peripheral TRMs. Surprisingly, however, the effect conferred by butyrate appeared to require prolonged exposure, as overnight exposure (~16h) did not significantly enhance efferocytosis efficiency (Fig. S2B). Additionally, the boost in efferocytosis capacity and efficiency provided by butyrate were maintained for up to 2d after withdrawal of butyrate *in vitro* (**Fig. 2D** and Fig. S2C).

**Fig. 2.**
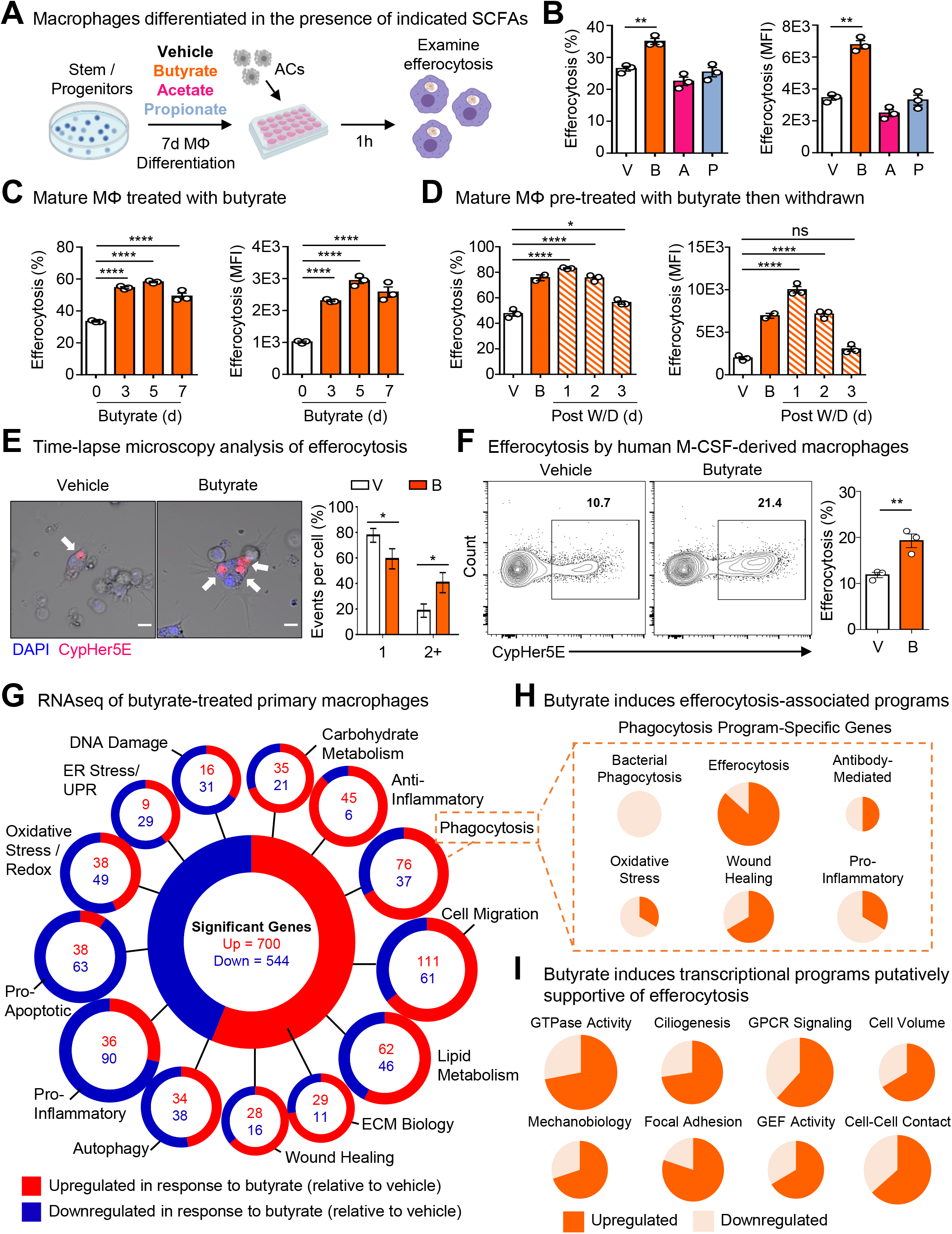
Butyrate boosts efferocytosis via induction of efferocytotic transcriptional programs. (A, B) <u>Exogenous butyrate, but not other short-chain fatty acids (SCFAs), boosts efferocytosis by primary macrophages *in vitro*</u>. (A) Schematic of experimental design. Bone marrow was differentiated into macrophages in the presence of butyrate, acetate, or propionate (all 1 mM) for 7d. Mature macrophages were subsequently incubated with apoptotic cells co-labeled with CypHer5E and CellTrace Yellow (CTY) at a 1:1 ratio for 1h. (B) Shown is summary data from flow cytometry analysis of efferocytosis efficiency (left, CypHer5E+ macrophages) and capacity (right, CTY median fluorescence intensity, MFI in macrophages) of experiments performed as described in (A). Data are from three independent experiments. \*\**p* < .01. (C) <u>Mature macrophages conditioned with butyrate exhibit enhanced efferocytosis *in vitro*</u>. Mature macrophages were conditioned with butyrate (1 mM) for 0-7d prior to efferocytosis assay. Conditioned macrophages were subsequently incubated with apoptotic cells co-labeled with CypHer5E and CTY at a 1:1 ratio for 1h. Shown is summary data from flow cytometry analysis of efferocytosis efficiency (left, CypHer5E+ macrophages) and capacity (right, CTY MFI in macrophages). Data are from three independent experiments. \*\*\*\**p* < .0001. (D) <u>The effect of butyrate on mature macrophages is reversible</u>. Mature macrophages were conditioned with butyrate (1 mM) for 3d followed by withdrawal of butyrate for the indicated time point (1-3d). Conditioned macrophages were subsequently incubated with apoptotic cells co-labeled with CypHer5E and CTY at a 1:1 ratio for 1h. Shown is summary data from flow cytometry analysis of efferocytosis efficiency (left, CypHer5E+ macrophages) and capacity (right, CTY MFI in macrophages). Data are from three independent experiments. \*\*\*\**p* < .0001. ns = not significant. (E) <u>Mature macrophages conditioned with butyrate engulf more apoptotic cells on a per-cell basis</u>. Time-lapse confocal microscopy analysis of macrophages conditioned as in (C) then cultured with CypHer5E-labeled apoptotic cells. Macrophages containing at least one CypHer5E puncta (white arrows) were analyzed for the number of apoptotic cell uptake (CypHer5E+) events. Data is shown as the fraction of the total number of CypHer5E+ events. Scale bar, 10μm. Data are from three independent experiments. \**p* < .05. (F) <u>Human macrophages conditioned with butyrate also exhibit enhanced efferocytosis *in vitro*</u>. Human monocyte-derived macrophages (HMDMs) were conditioned with vehicle or butyrate (1 mM; 3d) then incubated with CypHer5E-labeled apoptotic cells at a 1:1 ratio for 1h. Shown is summary data from flow cytometry analysis of efferocytosis efficiency (left, CypHer5E+ macrophages). Data are from three independent experiments. \*\**p* < .01. (G) <u>RNAseq analysis of mature macrophages conditioned with butyrate</u>. Mature macrophages were conditioned with vehicle or butyrate (1 mM) for 3d. Conditioned macrophages were subsequently lysed, and mRNA was extracted for downstream processing and sequencing. We observed significant changes in 1,244 genes in butyrate-conditioned macrophages relative to vehicle-conditioned macrophages. Differentially expressed genes were classified according to known or putative (based on family or sequence similarity) function. Transcripts were considered significant if they fell below a false discovery rate-adjusted *p* value < .05. Data are from three independent experiments. ECM, extracellular matrix. (H) <u>Butyrate induces efferocytosis-associated transcriptional programs in mature macrophages</u>. Transcripts that were categorized as ‘phagocytosis’ in (G) were subsequently analyzed for additional functional signatures. (I) <u>Butyrate induces transcriptional programs putatively supportive of efferocytosis in mature macrophages</u>. Transcripts that were significantly differentially regulated in response to butyrate conditioning as identified in (G) were subsequently analyzed for putative efferocytosis-associated functional programs.

Because TRMs are the primary professional phagocyte responsible for clearing millions of dead cells daily (*1*), the ability to internalize multiple apoptotic cells simultaneously is likely an essential ability (*53*). We therefore asked whether butyrate treatment of macrophages alters efferocytosis capacity on a per-cell basis. To assess this, we performed time-lapse confocal microscopy of vehicle- and butyrate-treated primary macrophages cultured with apoptotic cells. The majority of vehicle-treated macrophages (~80%) engulfed one apoptotic cell, with all vehicle-treated macrophages engulfing one or two apoptotic cells (**Fig. 2E**). Contrarily, a significantly higher fraction of butyrate-treated macrophages (~40%) engulfed two or more apoptotic cells, with several macrophages engulfing three or more apoptotic cells (**Fig. 2E**). Fascinatingly, butyrate-treated macrophages exhibited a pronounced change in morphology, exemplified by the extension of numerous filopodia (Fig. S2D). Such filopodial extensions are thought to correspond with a faster form of phagocytosis (*54*) and to be important for efficient sensing of cell death and phagocytosis by embedded tissue-resident macrophages (*54*, *55*). In support of the notion that butyrate conditions macrophages to be ‘better’ phagocytes, we also observed that butyrate-treated macrophages exhibit enhanced antibody-mediated phagocytosis compared to vehicle-treated controls (Fig. S2E). Importantly, we found that the benefit of butyrate extends to human macrophages, as butyrate treatment significantly enhanced efferocytosis by primary human monocyte-derived macrophages (**Fig. 2F**). Collectively, our findings suggest that butyrate informs macrophage efferocytosis potential.

In order to better understand why macrophages treated with butyrate are more efficient phagocytes, we performed bulk mRNA sequencing (RNAseq) of primary macrophages treated with butyrate or a vehicle control. Analysis of butyrate-treated macrophages revealed differential regulation of numerous cell biological function-associated transcriptional programs (**Fig. 2G** and Table S1). For example, butyrate-treated macrophages significantly upregulate both carbohydrate and lipid metabolism programs but downregulate both oxidative stress/redox biology and ER Stress/Unfolded Protein Response (UPR) programs (**Fig. 2G**). Importantly, several of these programs, such as wound healing, extracellular matrix (ECM) biology, and phagocytosis, are unique associations with butyrate treatment of primary macrophages, in contrast to monocyte-derived macrophages differentiated in the presence of butyrate (*56*). Broadly, our RNAseq analysis suggests that butyrate treatment of primary macrophages enforces core tissue-resident macrophage programs important for homeostatic function.

Given our finding that butyrate treatment enhances efferocytosis efficiency and capacity, we next explored in more detail the phagocytosis transcriptional program induced by butyrate treatment. The phagocytosis transcriptional program is a broad categorization that includes various homeostatic- and pro-inflammatory-associated programs. Strikingly, we observed that butyrate treatment significantly downregulates genes associated with bacterial phagocytosis, oxidative stress/redox biology, and pro-inflammatory signaling (**Fig. 2H**) within the phagocytosis program. Contrarily, we observed significant upregulation of genes associated with efferocytosis and wound healing (**Fig. 2H**) as well as cell adhesion, cell migration, and anti-inflammatory signaling genes (Fig. S2F) within the phagocytosis program. Efferocytosis consists of discrete steps that require distinct cell biological processes and signaling machinery (*57*). Interestingly, analysis of butyrate-treated macrophages revealed significant upregulation of transcriptional programs putatively involved in efferocytosis, such as GTPase activity, cell volume regulation, and focal adhesion (**Fig. 2I** and Fig. S2G). Taken together, our data suggest that butyrate induces transcriptional programs in macrophages that support core tissue macrophage functions, including efferocytosis.

### Exogenous butyrate rescues antibiotic-induced defects in peripheral efferocytosis

We next sought to determine if butyrate availability could account for the defective peripheral efferocytosis observed in antibiotic-treated mice. To test this, we took two independent *in vivo* approaches. First, we supplemented the drinking water of antibiotic-treated mice with butyrate to imitate the conventional route of access (**Fig. 3A**). Similar to our previous observations, efferocytosis was strikingly decreased in antibiotic-treated mice compared to control mice (**Fig. 3A**, compare blue and green bars). On the other hand, antibiotic-treated mice orally supplemented with butyrate exhibited significantly rescued efferocytosis capacity (**Fig. 3A**, compare green and orange bars). Second, we delivered butyrate directly to the peritoneal cavity, allowing us to test both the direct relevance of butyrate on pMac efferocytosis *in vivo* and to bypass potential confounds that might arise from dysbiosis in the intestines (**Fig. 3B**). Similar to oral supplementation, peritoneal delivery of butyrate significantly rescued pMac efferocytosis in antibiotic-treated mice (**Fig. 3B**, compare green and orange bars). Thus, intestinal microbiome-derived butyrate is essential for efferocytosis by peripheral TRMs.

**Fig. 3.**
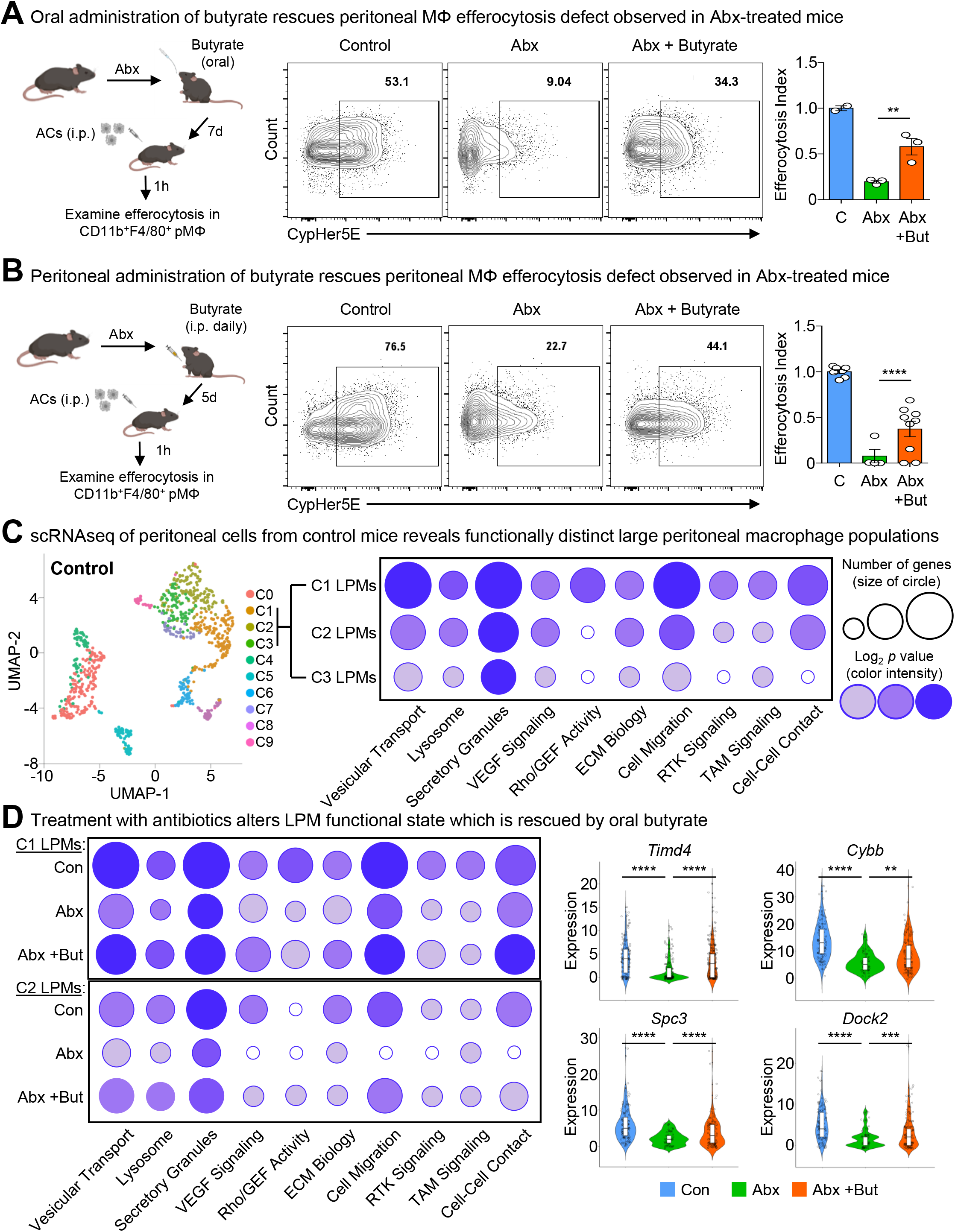
Exogenous butyrate rescues antibiotic-induced defects in peripheral efferocytosis. (A) <u>Oral administration of butyrate rescues efferocytosis capacity by peritoneal macrophages in antibiotic-treated mice *in vivo*</u>. (left). Schematic of experimental design. Mice were initially treated with antibiotics for 7d. Then, half of antibiotic-treated mice were supplemented with oral butyrate (150 mM) in the drinking water for 7d. Mice were subsequently injected intraperitoneally (i.p.) with CypHer5E-labeled apoptotic cells (ACs). After 1h, peritoneal CD11b^+^F4/80^+^ macrophages were analyzed for efferocytosis. Shown are representative flow cytometry (middle) and summary (right) plots of the rate of efferocytosis by CD11b^+^F4/80^+^ macrophages in control mice (n = 2), in mice treated with antibiotics (ABX, n = 3), and in mice with antibiotics supplemented with butyrate (ABX+But, n = 3). Statistical analysis was performed only between ABX and ABX+But mice. \*\**p* < .01. Data are from one independent experiment. (B) <u>Peritoneal delivery of butyrate rescues efferocytosis capacity by peritoneal macrophages in antibiotic-treated mice *in vivo*</u>. (left). Schematic of experimental design. Mice were initially treated with antibiotics for 7d. Then, half of antibiotic-treated mice were supplemented with daily i.p. injection of butyrate (40 mM) for 5d. Mice were subsequently injected i.p. with CypHer5E-labeled ACs. After 1h, peritoneal CD11b^+^F4/80^+^ macrophages were analyzed for efferocytosis. Shown are representative flow cytometry (middle) and summary (right) plots of the rate of efferocytosis by CD11b^+^F4/80^+^ macrophages in control mice (n = 7), mice treated with antibiotics (ABX, n = 4), and in mice treated with antibiotics supplemented with i.p. butyrate (ABX+But, n = 9). Data are from two independent experiments. \*\*\*\**p* < .0001. (C, D) <u>Single cell RNA sequencing analysis of peritoneal cells from control mice, mice treated with antibiotics, and mice treated with antibiotics and oral butyrate</u>. (C) Peritoneal cells were isolated, labeled with barcoded MHC Class I-specific antibodies, and processed for scRNAseq. Shown is the dimension reduction analysis via Uniform Manifold Approximation and Projection (UMAP) of control peritoneal cells (left). Individual cell expression profiles were cross-referenced with cell type-specific transcription profiles curated by the Immunological Genomes project, allowing for the identification of specific cell types including large peritoneal macrophages (LPMs; see also Fig. S3 and Fig. S4). (right) Functional analysis of differentially expressed genes extrapolated from the three main LPM clusters (C1-C3). The circle size was determined by applying a scaling factor to the number of genes involved in the program. The upper bound features 120-150 differentially expressed genes, which is scaled to 0.35” whereas the lower bound features 0-19 differentially expressed genes, which is scaled to 0.16”. The color intensity was determined by applying a scaling factor to the log_2_ *p* value. The upper bound (100% intensity) features log_2_ *p* values greater than 100 whereas the lower bound (0% intensity) features log_2_ *p* values less than 15. (D) Functional programs identified as enriched in C1 and C2 LPMs from control mice were subsequently analyzed in mice treated with antibiotics and mice treated with antibiotics supplemented with oral butyrate. Shown are the summary plots of functional programs (left) and representative genes (right).

Although we were able to rescue peripheral TRM efferocytosis by providing butyrate locally, it remains possible that microbiota-derived butyrate does not act directly on peripheral TRMs. To further explore this idea, we performed single cell RNAseq (scRNAseq) of peritoneal cells isolated from control mice, antibiotic-treated mice, and antibiotic-treated mice supplemented with oral butyrate (**Fig. 3C**). We initially performed an unbiased analysis of the peritoneal immune compartment to determine if antibiotic treatment was affecting the peritoneum more broadly. However, uniform manifold approximation and projection (UMAP) analysis revealed a consistent presence of key immune cell subsets (*58*), including B1 B Cells (*Ms4a1*+), T cells (*Il7r*+), Small Peritoneal Macrophages (SPMs; *Clec4b1*+), and Large Peritoneal Macrophages (LPMs; *Itgam*+, *Adgre1*+), across all three conditions (Fig. S3 and S4). Furthermore, by cross-referencing our expression data with immune subset RNAseq available via the Immunological Genome project (*59*), we identified four overlapping but distinct LPM populations in untreated mice, characterized by transcriptional expression of F4/80 (*Adgre1*), CD11b (*Itgam*), FCER1G (*Fcer1g*), and LysM (*Lyz2*). The predominant LPM population in untreated mice (Cluster 1) most closely resembled the canonical phagocytic peritoneal TRM, featuring robust expression of *Adgre1* and *Itgam* as well as significant expression of efferocytosis receptors *Timd4*, *Mertk*, and *Tyrobp* (Fig. S3 and S4; Table S2). This population also accounted for the majority of individual LPMs. The second LPM population (Cluster 2) also featured significant *Adgre1* and *Itgam* expression as well as increased *Timd4* and *Tyrobp* expression but did not exhibit enhanced *Mertk* expression (Fig. S3 and S4; Table S2). The third (Cluster 3) and fourth (Cluster 7) LPM populations featured enrichment of genes uniquely found in LPM RNAseq from ImmGen but were instead predominantly characterized by enrichment of other previously identified LPM genes, such as *Fcer1g* (Cluster 3) and *Lyz2* (Cluster 7) and lacked canonical efferocytosis receptors *Timd4* and *Mertk* (Fig. S3 and S4; Table S2). Nonetheless, identification of LPM clusters allowed us to perform downstream analysis of the impact of antibiotic treatment and butyrate rescue on LPM functional programs.

Further analysis of LPMs from control mice revealed that the peritoneum consists of two populations of LPMs enriched for functional programs putatively supportive of efferocytosis: Cluster 1 and Cluster 2 LPMs (**Fig. 3C** and Table S3). Specifically, we observed enrichment of programs, including putative efferocytosis programs, induced in macrophages treated with butyrate *in vitro*, including vesicular transport, lysosome biology, Rho/GEF activity, ECM biology, cell migration, receptor tyrosine kinase (RTK) signaling, and cell-cell contact/cell adhesion (**Fig. 3C** and Table S3). Strikingly, both Cluster 1 and Cluster2 LPMs from antibiotic-treated mice exhibited significant decreases across analyzed functional programs, especially lysosome biology, Rho/GEF activity, and RTK signaling (**Fig. 3D**). Furthermore, amongst the identified LPM populations in antibiotic-treated mice, there were fewer cells expressing core efferocytosis transcripts and the expression level of core transcripts was lower (e.g., *Timd4*; **Fig. 3D** and Table S3). On the other hand, both Cluster 1 and Cluster 2 LPMs from antibiotic-treated mice who received oral butyrate supplementation exhibited a significant rescue of analyzed functional programs, including putative efferocytosis programs (**Fig. 3D**). Our data indicate that microbiome-derived butyrate supports efficient efferocytosis by peripheral tissue-resident macrophages through direct regulation of efferocytosis-associated transcriptional programs.

### Butyrate boosts efferocytosis via histone acetylation, not cognate GPCR signaling

Short-chain fatty acids (SCFAs) exert effects on cellular function through at least two routes: via extracellular ligation of G protein-coupled receptors (GPCRs) or via transport-mediated uptake and subsequent regulation of redox metabolism and transcription (*60*). To date, the majority of research suggests that microbiome-derived butyrate affects intestinal immune cell function through GPCR signaling and inhibition of histone deacetylase activity (*61*, *62*). RNAseq analysis revealed that pMacs, at least transcriptionally, appear to express the cognate butyrate GPCR GPR109a (*Hcar2*), with negligible expression of the other known SCFA GPCRs GPR41 (*Ffar3*) or GPR43 (*FFar2*) (**Fig. 4A**), consistent with publicly available data from BioGPS and ImmGen databases (*59*, *63*). Accordingly, we first investigated whether GPCR signaling could account for the enhanced efferocytosis promoted by butyrate (**Fig. 4B**). Contrary to our hypothesis, macrophages lacking GPR109a exhibited enhanced efferocytosis in response to butyrate *in vitro* (**Fig. 4C**, compare white and pink bars). Given previous studies have reported macrophage function relying on GPR43 (*64*), we also tested whether GPR43 is required for butyrate-mediated enhanced efferocytosis. Similar to our results with GPR109a-deficient macrophages, GPR43-deficient macrophages treated with butyrate exhibited significantly enhanced efferocytosis capacity *in vitro* (**Fig. 4C**, compare white and green bars). Additionally, culturing macrophages with the GPR109a agonist niacin failed to enhance efferocytosis capacity (**Fig. 4D**). Although the *in vitro* methods we use to study macrophage efferocytosis are generally good approximations of TRM function, there remain unique factors in the tissue environment that may inform TRM biology (*3*). To further assess if GPR109a and GPR43 are required for peripheral macrophage efferocytosis, we performed efferocytosis assays in GPR109a- and GPR43-deficient mice. Consistent with our *in vitro* results, we observed no significant differences between wildtype and GPR109a-deficient mice (**Fig. 4E** and Fig. S5A) or between wildtype and GPR43-deficient mice (Fig. S5B and S5C). Taken together, our results suggest that butyrate signaling via known cognate GPCRs is dispensable for efficient efferocytosis.

**Fig. 4.**
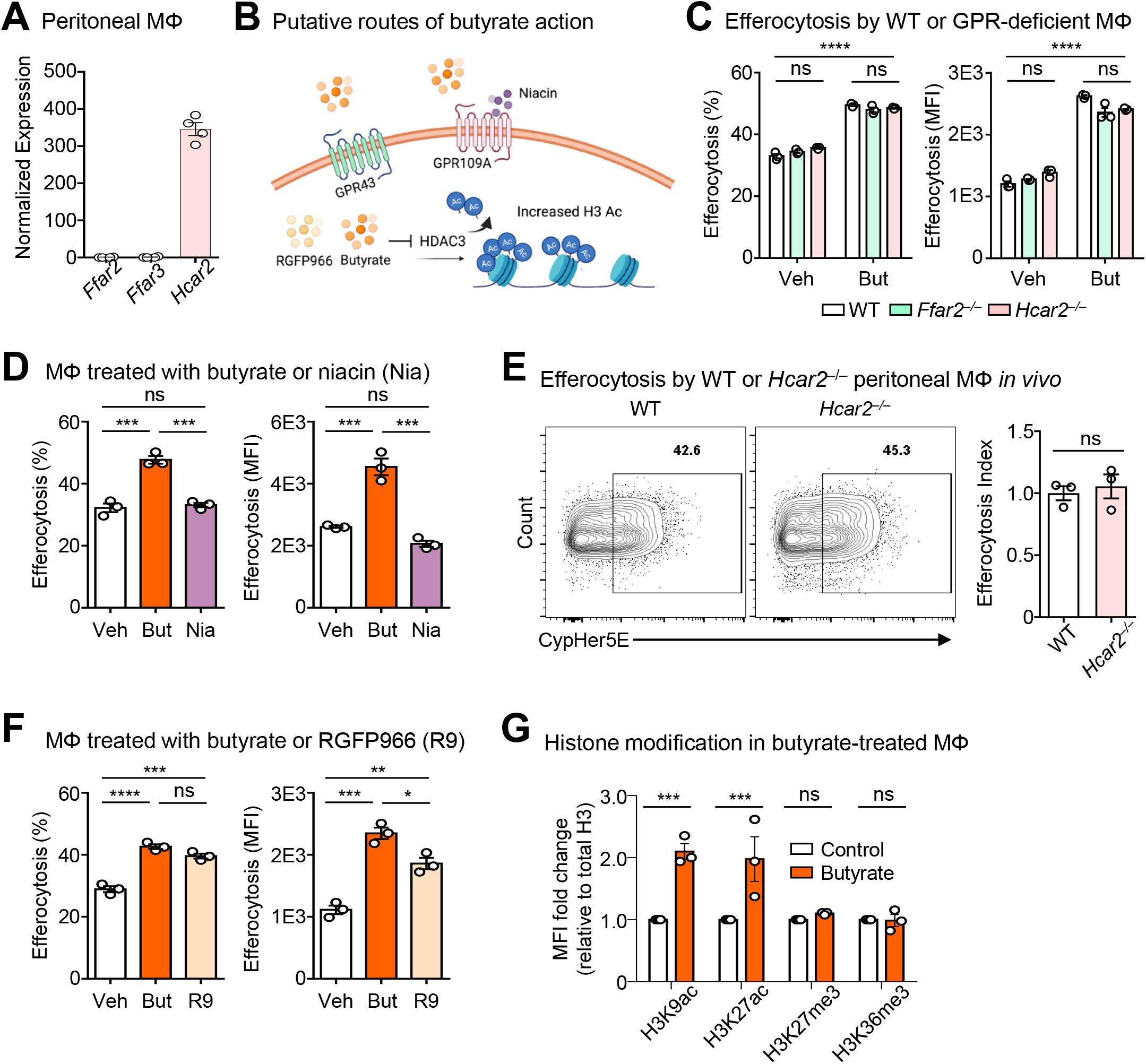
Butyrate boosts efferocytosis via histone acetylation, not cognate GPCR signaling. (A) <u>Naïve peritoneal macrophages specifically express *Hcar2*</u>. Analysis of *Ffar2* (protein name: GPR43), *Ffar2* (protein name: GPR42), and *Hcar2* (protein name: GPR109a) normalized gene expression in bulk RNAseq of isolated peritoneal macrophages from male, C57BL/6 mice (n=4). (B) <u>Putative routes of butyrate action</u>. Shown is a schematic illustrating experiments performed to interrogate the mechanism of butyrate action, testing specifically if butyrate acts via cognate GPCR or inhibition of HDAC3. (C) <u>Efferocytosis by primary macrophages does not require GPR109a or GPR43 *in vitro*</u>. Macrophages were derived from wildtype (WT; white), *Ffar2*^−/−^ (green) and *Hcar2*^−/−^ (pink) mice and conditioned with vehicle or butyrate (1 mM) for 3d. Conditioned macrophages were subsequently incubated with CypHer5E and CellTrace Yellow (CTY)-labeled apoptotic cells (ACs) at a 1:1 ratio for 1h. Shown is summary data from flow cytometry analysis of efferocytosis efficiency (left, CypHer5E+ macrophages) and capacity (right, CTY MFI in macrophages). Data are from three independent experiments. \*\*\*\**p* < .0001. ns = not significant. (D) <u>Induction of GPR109a by niacin does not boost efferocytosis by primary macrophages *in vitro*</u>. Mature macrophages were conditioned with vehicle, butyrate (1 mM), or the GPCR109a activator niacin (500 μM) for 3d prior to efferocytosis assay. Conditioned macrophages were subsequently incubated with ACs co-labeled with CypHer5E and CTY at a 1:1 ratio for 1h. Shown is summary data from flow cytometry analysis of efferocytosis efficiency (left, CypHer5E+ macrophages) and capacity (right, CTY MFI in macrophages). Data are from three independent experiments. \*\*\**p* < .001. ns = not significant. (E) <u>Efferocytosis by peritoneal macrophages does not depend on GPR109a *in vivo*</u>. *Hcar2*^−/−^ (n=3) mice were injected intraperitoneally (i.p.) with CypHer5E-labeled ACs. After 1h, peritoneal CD11b^+^F4/80^+^ macrophages were analyzed for efferocytosis. Shown are representative flow cytometry (left) and summary (right) plots of the rate of efferocytosis by CD11b^+^F4/80^+^ macrophages. ns = not significant. Data are from two independent experiments. (F) <u>Inhibition of HDAC3 boosts efferocytosis by primary macrophages *in vitro*</u>. Mature macrophages were conditioned with vehicle, butyrate (1 mM), or the HDAC3 inhibitor RGFP966 (R9; 20 μM) for 3d prior to efferocytosis assay. Conditioned macrophages were subsequently incubated with ACs co-labeled with CypHer5E and CTY at a 1:1 ratio for 1h. Shown is summary data from flow cytometry analysis of efferocytosis efficiency (left, CypHer5E+ macrophages) and capacity (right, CTY MFI in macrophages). Data are from three independent experiments. \**p* < .05; \*\**p* < .01; \*\*\**p* < .001. \*\*\*\**p* < .0001. ns = not significant. (G) <u>Exogenous butyrate induces increased H3K9 and H3K27 acetylation in primary macrophages *in vitro*</u>. Primary macrophages were conditioned as in (F), then subsequently analyzed for selected histone modifications. Shown are summary plots of the MFI fold change observed in butyrate-treated macrophages compared to control macrophages (see also Fig. S5D for representative raw data plots). All samples were normalized to total H3. Data are from three independent experiments \*\*\**p* < .001. ns = not significant.

Butyrate also functions intracellularly by inhibiting histone deacetylases (HDACs), which leads to enhanced and sustained histone acetylation and subsequent regulation of gene expression (*65*). We next investigated whether direct HDAC inhibition impacts efferocytosis capacity. To this end, we treated macrophages with RGFP966, a specific inhibitor of the known butyrate target HDAC3 (**Fig. 4B**) (*56*, *65*). Inhibition of HDAC3 significantly enhanced efferocytosis efficiency and capacity to a level similar to butyrate-conditioned macrophages (**Fig. 4F**), suggesting that butyrate supports peripheral efferocytosis through inhibition of HDAC3. Additionally, treatment with butyrate increased the acetylation of both lysines 9 and 27 on histone 3 (H3K9ac and H3K27ac) but had no effect on tri-methylation of lysines 27 and 36 (H3K27me3 and H3K36me3) (**Fig. 4G** and Fig. S5D), further supporting the notion that butyrate supports peripheral efferocytosis via cell-intrinsic transcription regulation via histone deacetylation. Our results suggest that microbiome-derived butyrate functions to transcriptionally ‘prepare’ peripheral macrophages for homeostatic functions, including efferocytosis.

### Treatment with antibiotics induces prolonged peripheral efferocytosis defects

Dysbiosis induced by antibiotics use may have lasting effects long after completion of the course. Previous work has demonstrated that intestinal microbiome reconstitution takes weeks to months, with certain bacterial groups failing to return years after antibiotic withdraw (*66*–*70*). These findings coincide with epidemiological observations of an association between antibiotics use and non-intestinal autoimmunity (e.g., rheumatoid arthritis (*31*, *71*)). Efferocytosis, on the other hand, occurs in every major tissue and prolonged perturbations of efferocytosis can lead to failed wound healing and repair, with chronic failure thought to lead to autoimmunity and chronic inflammatory disease (*2*, *4*, *5*, *72*). Given these clinical and epidemiological observations, we queried whether treatment with antibiotics induced disruption of peripheral efferocytosis beyond the end of the course (**Fig. 5A**). Consistent with our previous experiments, we observed decreased efferocytosis by peritoneal macrophages (pMacs) in antibiotic-treated mice one day post-antibiotic withdrawal (**Fig. 5B**, left graph). This deficiency corresponded with the hallmark enlarged, dark-colored cecum found in mice with dysbiosis (**Fig. 5B**, left image). Strikingly, peripheral efferocytosis remained significantly impaired beyond two weeks post-antibiotic withdrawal, despite the apparent reversal of dysbiosis (**Fig. 5B**, center). The peripheral efferocytosis deficiency was subsequently reversed by three weeks post-antibiotic withdrawal (**Fig. 5B**, far right), further supporting the notion that peripheral efferocytosis is informed by microbiota-derived signals, including butyrate.

**Fig. 5.**
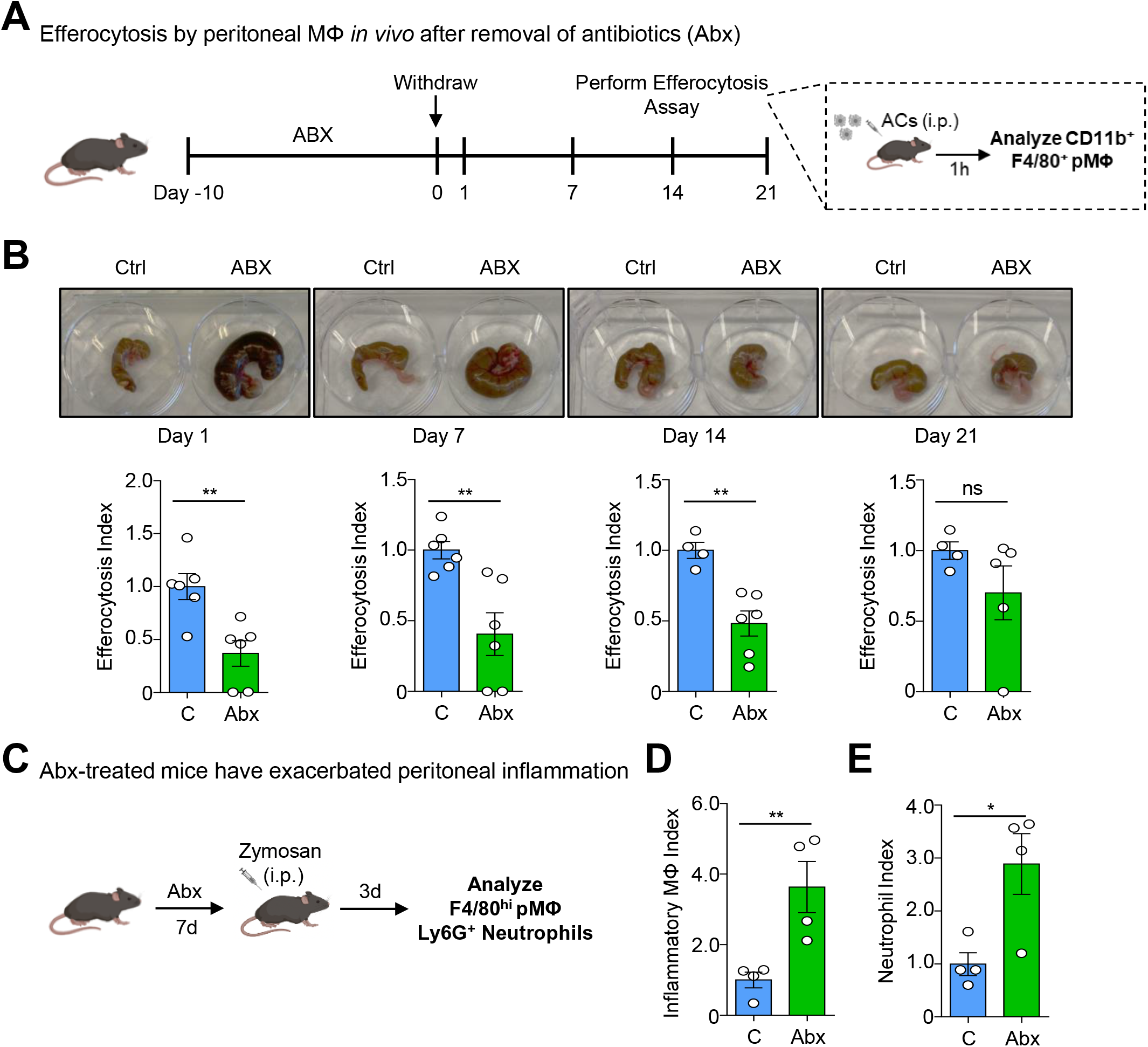
Treatment with antibiotics induces prolonged peripheral efferocytosis defects. (A, B) <u>Treatment with antibiotics induces a prolonged defect in peripheral efferocytosis *in vivo*</u>. (A) Schematic of experimental design. Mice were treated with antibiotics (ABX) in drinking water for 10d followed by withdrawal of antibiotics. On indicated days post-antibiotics withdrawal, mice were injected intraperitoneally (i.p.) with CypHer5E-labeled apoptotic cells. After 1h, peritoneal CD11b^+^F4/80^+^ macrophages were analyzed for efferocytosis. (B) Shown are representative images of ceca (top) and summary plots of the rate of efferocytosis by CD11b^+^F4/80^+^ macrophages (bottom). Data are from day 1 (control n=6, ABX n=6), day 7 (control n=6, ABX=6), day 14 (control n=4, ABX=6), and day 21 (control=4, ABX=4) post-antibiotics withdrawal, across three independent experiments. \*\**p* < .01. ns = not significant. (C-E) <u>Mice treated with antibiotics are susceptible to peritoneal injury</u> (C) Schematic of experimental design. Mice were treated with antibiotics in the drinking water for 7d prior to i.p. injection of zymosan (0.1 mg). After 3d, when inflammatory cell infiltration has resolved in naïve adult mice, peritoneal lavage was collected and analyzed by flow cytometry. (D) Shown are summary plots of inflammatory macrophages (F4/80^high^) and (E) apoptotic neutrophils (AnnexinV^+^CD11b^+^Ly6G^+^), normalized to average values in control mice. Data are from two independent experiments, with two mice per condition (n=4 per group). \**p* < .05; \*\**p* < .01.

Lastly, we sought to determine whether treatment with antibiotics affects susceptibility to an efferocytosis-dependent sterile inflammation resolution model. To this end, we utilized the zymosan-induced peritoneal damage model which depends on peritoneal macrophage clearance of apoptotic neutrophils to resolve inflammation (*73*). Indeed, mice treated with antibiotics were less capable of resolving inflammation upon challenge (**Fig. 5C**), including prolonged presence of pro-inflammatory macrophages (**Fig. 5D** and Fig. S6) and neutrophilia (**Fig. 5E** and Fig. S6). Consequently, antibiotic treatment leads to prolonged deficiency in homeostatic peripheral efferocytosis beyond completion of the antibiotics course and leaves the host susceptible to peripheral inflammatory injury.

## Discussion

The ability of phagocytes to clear apoptotic cells is a key facet of tissue health, especially in the context of sterile injury. Indeed, organisms that exhibit perturbations of efferocytosis machinery often present with inflammatory sequalae which, in mammals, can result in the development of inflammatory disease. How this ability is established and maintained in tissue-resident phagocytes (such as tissue-resident macrophages) and whether signals arising from distal sites (such as the intestinal tract) contribute remains poorly understood. Here, across multiple lines of investigation, we made the surprising discovery that the ability of peripheral phagocytes to perform efferocytosis is informed by signals arising from the intestinal microbiome. In particular, we found that mice treated with broad spectrum antibiotics exhibit significant efferocytosis defects in both the peritoneum and the thymus. This defect was attributable to the absence of the intestinal microbiome, as (1) germ-free mice also exhibited a significant decrease in peripheral phagocyte efferocytosis, and (2) antibiotic treatment-induced efferocytosis defects were rescued via fecal microbiota transplantation. Mechanistically, we discovered that the short-chain fatty acid butyrate is an essential distal signal capable of rescuing peripheral efferocytosis defects caused by antibiotics treatment. Through a combination of cell biological, bulk and single cell RNA sequencing, and *in vivo* animal approaches, we show that butyrate acts directly on peripheral macrophages, the main professional tissue-resident phagocyte, to support efferocytosis. Specifically, mouse and human macrophages conditioned with butyrate exhibited enhanced efferocytosis efficiency and capacity. Additionally, both oral and intraperitoneal administration of butyrate rescued the efferocytosis defect observed in peritoneal macrophages of antibiotic-treated mice *in vivo*. We found that butyrate supports macrophage efferocytosis via inhibition of HDAC3, not interaction with its cognate GPCRs, inducing transcriptional programs that are involved in or supportive of efferocytosis. Importantly, we observed that the defect in peripheral efferocytosis persists beyond antibiotic withdrawal and that mice treated with antibiotics had exacerbated inflammation in an efferocytosis-dependent sterile inflammation resolution model. Our work provides an additional possible link between the clinical observation that patients receiving broad-spectrum antibiotics are susceptible to the development of inflammatory disease and autoimmunity in non-intestinal tissues (*30*, *31*, *71*, *74*–*79*). Taken together, we propose that tissue-resident phagocytes rely on integration of distal molecular signals to establish and maintain their ability to perform efferocytosis.

The importance of butyrate on host immune cell function in the intestines is well-established (*20*, *21*), though its importance for intestinal macrophage function is less straightforward. For instance, previous work suggests that butyrate decreases responsiveness to the Gram-positive bacteria membrane component lipopolysaccharide, in particular, production of the pro-inflammatory mediators nitric oxide, IL-6, and IL-12 (*80*, *81*). However, a recent study suggested that butyrate, despite suppressing inflammatory cytokine production, programs monocyte-derived macrophages into effective antimicrobial macrophages including increased production of antimicrobial products in response to bacterial infection *in vitro* and in the intestines *in vivo* (*56*), consistent with the finding that macrophages treated with both IL-4 and butyrate efficiently internalize and kill bacteria (*81*). To our knowledge, our work is the first to examine the role of butyrate on macrophage identity and function outside of the intestines. Our findings complement this previous work by demonstrating that bacterial products regulate essential macrophage functions in non-barrier tissues. However, our finding that butyrate downregulates programs putatively involved in bacterial phagocytosis in mature macrophages raises the interesting possibility that the continuous replenishment of intestinal macrophages from blood monocytes (*82*) serves to reinforce the presence of semi-tolerogenic intestinal macrophages. Contrarily, tissue-resident macrophages, such as peritoneal LPMs, seed tissues during embryogenesis and are not exposed to bacteria until the host is faced with an infection. In these peripheral tissues, we propose that butyrate and possibly other microbial-derived signals, act specifically to enforce macrophage function essential to a given tissue.

Our work raises several interesting questions. For instance, we observed that exogenous butyrate, regardless of the route of supplementation, resulted in an approximate 50% rescue of efferocytosis by peritoneal macrophages. What additional intestinal microbiome-derived signals support peripheral efferocytosis? Although we report that efferocytosis is impaired in antibiotic-treated mice in both the peritoneum and the thymus, are other tissue-resident macrophage-specific clearance processes, such as the removal of mitochondria in the heart (*83*) or surfactant in the lung (*84*), also affected? Relatedly, most tissues have both professional and non-professional phagocytes that perform unique tissue-specific clearance activities (*85*). Do distal signals affect professional and non-professional phagocytes differently? Finally, do host tissues generate molecular signals that act distally to inform efferocytosis in other tissues? Nevertheless, the current study provides a roadmap to further our understanding of how tissue-resident phagocytes, in particular tissue-resident macrophages, are programmed to perform efferocytosis and maintain tissue homeostasis. The data presented here advance the concept that metabolites produced by commensal microbes act peripherally to enforce efficient dead cell clearance by tissue-resident phagocytes and may help to explain the epidemiological observation that patients treated with antibiotics exhibit increased incidence of inflammatory disease and autoimmunity.

## Materials and Methods

### Animal studies

Wild-type C57BL/6J (Stock No. 000664) and J:DO (Stock No. 009376) were purchased from The Jackson Laboratory. *Gpr43*^−/−^ and *Gpr109a*^−/−^ mice were provided by M.v.d. Brink. Germ-free mice were provided by A. Rudensky, maintained in flexible isolators, and routinely checked for bacteria and fungi by PCR of fecal DNA samples for bacterial 16S and fungal 18S, respectively. All mice were housed at the Research Animal Resource Center for MSKCC (specific-pathogen free) and Weill Cornell Medicine (germ-free) with 12-hour light/dark cycles. Mice were housed in ambient conditions and *ad libitum* access to water and food. All studies were conducted under protocol number 19-07-012 and approved by the Sloan Kettering Institute (SKI) Institutional Animal Care and Use Committee (IACUC).

### Reagents

The reagents used in this work were as indicated: CypHer5E (GE Life Sciences, PA15401); CellTrace Yellow (Thermo Fisher, C34567); Hoechst (Thermo Fisher, H3570); sodium butyrate (Sigma, 303410, 1 mM); sodium acetate (Sigma, S2889, 1 mM); sodium propionate (Sigma, P1880, 1 mM); RGFP966 (Selleck Chemicals, S7229, 20 μM); niacin (Sigma, N4126, 500 μM).

### Bone marrow-derived macrophages

BMDMs were generated by culturing mouse bone marrow cells in alpha-MEM (Fisher Scientific, MT15012CV) containing 10% (vol/vol) heat-inactivated FBS (Sigma), 10% (vol/vol) L929 cell-conditioned medium, 100 U/ml penicillin, 100 μg/ml streptomycin, and L-glutamine 37 °C and 5% CO_2_ for 7 days. BMDMs were seeded at indicated densities and allowed to rest overnight prior experiments.

### Human monocyte-derived macrophages

Peripheral blood mononuclear cells (PBMC) derived from healthy blood donors were obtained by gradient separation using 1-Step Polymorphs (Accurate Chemical, AN221725). CD14^+^ monocytes were isolated with the Mojosort human CD14 nanobeads (BioLegend, 480093). 5×10^6^ monocytes were seeded and differentiated into macrophages for 5 days in RPMI-1640 (Corning, 17-105-CV) supplemented with 10% (vol/vol) heat-inactivated FBS, 100 U/ml penicillin, 100 μg/ml streptomycin, L-glutamine, and 100 ng/ml recombinant human M-CSF (PeproTech, 300-25) at 37 °C and 5% CO_2_. HMDMs were seeded at indicated densities and allowed to rest overnight prior experiments.

### Antibiotic treatment

Mice were given a cocktail of antibiotics in their drinking water for at least 7 days prior experiments, unless otherwise stated. Antibiotic cocktail consisted of 1 g/L ampicillin (Cayman Chemical, 14417); 1 g/L kanamycin (K4000); 0.5 g/L metronidazole (Sigma, M3761); 0.5 g/L vancomycin (BioVision, B1507).

### Fecal microbiota transplant

About 6-8 stool pellets from SPF mice were collected into sterile micro tubes containing micro beads (Sarsted, 72.693.005), resuspended in sterile PBS, and vortexed until stool pellets were dissolved in solution. Fecal homogenates were centrifuged at max speed, and 200 μL were administered to mice through oral gavage. Mice were used for experiments 7 days after microbiota reconstitution.

### In vitro efferocytosis

Apoptosis was induced in human Jurkat T cells cultured in RPMI supplemented with 5% FBS by 150 mJ/cm ultraviolet C irradiation. Jurkat cells were incubated at 37 °C and 5% CO_2_ for 4 hours. For antibody-mediated phagocytosis, Jurkat cells were labelled with 25 μg/ml anti-human CD3 antibody (BioLegend, 317304, clone OKT3) and 3 μg/ml recombinant annexin V (BioLegend, 640901) for 1 hour at 4 °C. After, Jurkat cells were stained with the pH sensitive dye CypHer5E or CellTrace Yellow according to the manufacturer’s instructions before efferocytosis assay. After staining, Jurkat cells were resuspended in BMDM media. 1 × 10^5^ BMDMs were seeded in 24-well plates and stained with 2 μg/ml Hoechst to discriminate phagocyte from target. BMDMs were incubated with target cells at a 1:1 phagocyte:target ratio for the indicated times. Cells were scrapped off wells and phagocytes (Hoechst^+^) were assayed on an Attune NxT cytometer (Thermo Scientific). Samples were analyzed with FlowJo v10.8.1 (BD).

### In vivo efferocytosis

Mice were intraperitoneally injected with 1 × 10^6^ CypHer5E-labelled apoptotic Jurkat cells and euthanized 1-hour post-injection. Peritoneal lavage was collected in 8 ml cold PBS. Collected lavage was spun down and cells were blocked with CD16/32 prior being stained with CD11b eFluor 450 (eBioscience, 48-0112-82, clone M1/70, 1:200) and F4/80 PE (eBioscience, 12-4801-82, clone BM8, 1:400) for 30 min at 4 °C. Efferocytosis was analyzed by measuring CypHer5E^+^ events within the population of CD11b^+^F4/80^hi^ peritoneal macrophages in the flow cytometer. In separate *in vivo* efferocytosis experiments, six- to eight-week-old mice were injected i.p. with 300μl PBS containing 250 μg dexamethasone (50μM; Sigma). 6h post-injection, thymi were harvested from mice and the numbers of annexin V/7-AAD double positive cells (secondarily necrotic) were assessed by FACS.

### Zymosan-induced peritonitis

Mice were intraperitoneally injected with 0.1 mg of zymosan (Invivogen). Peritoneal lavage was collected in 8 ml cold PBS 3 days post-treatment. Collected lavage was spun down and cells were blocked with CD16/32 prior being stained with Ly6G PE-Cy7 (Tonbo Biosciences, 60-1276-U100, clone 1A8, 1:200) and F4/80 APC (BioLegend, 123116, clone BM8, 1:200) for 30 min at 4 °C and assessed by FACS.

### Butyrate treatment

BMDMs were treated with butyrate during or after differentiation for specific experiments. For *in vivo* local treatment, mice were treated intraperitoneally daily with 40 mM butyrate for the indicated times prior experiments. For oral treatment, mice that have been on antibiotic treatment were placed on drinking water with antibiotics supplemented with 100 mM butyrate for at least 10 days prior experiments.

### Histone acetylation and methylation analysis

For histone analysis, BMDMs were scrapped off wells after indicated treatments. Cells were fixed and permeabilized with the Foxp3 transcription factor staining kit (eBioscience, 00-5523-00) prior intracellular staining with the indicated antibodies: H3K9ac (Cell Signaling, 9649, 1:3000), H3K27ac (Cell Signaling, 8173, 1:2000), H3K27me3 (Cell Signaling, 9733, 1:2500), H3K36me3 (Active Motif, 61021, 1:2000), and total H3 (Active Motif, 39763, 1:500). Samples were then stained with anti-mouse (Invitrogen, A32728, 1:500) or anti-rabbit (Invitrogen, A32733, 1:500) Alexa Fluor 647 secondary antibodies, followed by flow cytometry analysis. Histone acetylation and methylation MFI was normalized to total H3 MFI from matching samples.

### Bulk RNA sequencing

Total RNA was isolated from 2 × 10^5^ cells untreated or butyrate-conditioned BMDMs using the NucleoSpin RNA extraction kit (Macherey-Nagel, 740902.50) according to the manufacturer’s instructions. An mRNA library was prepared by poly(A) enrichment using the Illumina TrueSeq platform. Samples were sequenced using the Illumina PE150 platform.

### Single-cell RNA sequencing and data analysis

Peritoneal lavage cells were collected as described above. Cells from different mice were stained with the following TotalSeq-A hashtag antibodies (all from BioLegend): Hashtag 1 (A0301), Hashtag 2 (A0302), Hashtag 3 (A0303), Hashtag 4 (A0304), Hashtag 5 (A0305), Hashtag 6 (A0306). Library preparation and single-cell RNA sequencing were performed by the Single Cell Research Initiative at SKI using 10X genomics Chromium Single Cell 3’ Library & Gel bead kit V3.1 according to the manufacturer’s instructions.

### Statistics and Reproducibility

Data were analyzed using GraphPad Prism 7, SPSS v.22, and R v.4.2.0. We determined significance using one of the following: unpaired two-tailed Student’s t-test, nonparametric Mann–Whitney U-test, one-way or two-way ANOVA, or Fisher’s exact test. R v.4.2.0 was used for graphical and statistical analyses and the R package DESeq2 was used for differential gene expression analysis of bulk RNAseq. Analysis of single-cell RNAseq data was performed using Python v.3.8 and R v.4.2.0. Specifically, the anaconda packages numpy, pandas, matplotlib, and scanpy as well as the R implementation of the standard Seurat v.4.0 pipeline were used in the course of data normalization, dimensionality reduction, clustering, and visualization. Gene functions were determined an approach described previously (*86*). In parallel, significantly differentially expressed genes were analyzed using the Molecular Signatures Database (MSigDB). All biologically independent samples are included in the statistical and graphical analyses featured in the manuscript and no data was excluded. Sample sizes were not predetermined using statistical methods. The code used in this manuscript is available upon reasonable request.

## Notes

### Competing Interest Statement

The authors have declared no competing interest.

## References

1. R. Sender, R. Milo, The distribution of cellular turnover in the human body. Nat. Med. 2021 271. 27, 45–48 (2021).

2. A. C. Doran, A. Yurdagul, I. Tabas, Efferocytosis in health and disease. Nat. Rev. Immunol. 20, 254–267 (2020).

3. G. Zago, P. H. V. Saavedra, K. R. Keshari, J. S. A. Perry, Immunometabolism of Tissue-Resident Macrophages – An Appraisal of the Current Knowledge and Cutting-Edge Methods and Technologies. Front. Immunol. 12, 1406 (2021).

4. S. Morioka, C. Maueröder, K. S. Ravichandran, Living on the Edge: Efferocytosis at the Interface of Homeostasis and Pathology. Immunity (2019),, doi:10.1016/j.immuni.2019.04.018.

5. E. Boada-Romero, J. Martinez, B. L. Heckmann, D. R. Green, The clearance of dead cells by efferocytosis. Nat. Rev. Mol. Cell Biol. 21, 398–414 (2020).

6. J. V. Patankar, C. Becker, Cell death in the gut epithelium and implications for chronic inflammation. Nat. Rev. Gastroenterol. Hepatol. 17 (2020), pp. 543–556.

7. Z. Szondy, É. Garabuczi, G. Joós, G. J. Tsay, Z. Sarang, Impaired clearance of apoptotic cells in chronic inflammatory diseases: Therapeutic implications. Front. Immunol. (2014),, doi:10.3389/fimmu.2014.00354.

8. A. Mahajan, M. Herrmann, L. E. Muñoz, Clearance deficiency and cell death pathways: A model for the pathogenesis of SLE. Front. Immunol. (2016),, doi:10.3389/fimmu.2016.00035.

9. M. C. Greenlee-Wacker, Clearance of apoptotic neutrophils and resolution of inflammation. Immunol. Rev. (2016),, doi:10.1111/imr.12453.

10. S. Arandjelovic, K. S. Ravichandran, Phagocytosis of apoptotic cells in homeostasis. Nat. Immunol. (2015),, doi:10.1038/ni.3253.

11. C. Z. Han, K. S. Ravichandran, Metabolic connections during apoptotic cell engulfment. Cell (2011),, doi:10.1016/j.cell.2011.12.006.

12. J. D. Proto, A. C. Doran, G. Gusarova, A. Yurdagul, E. Sozen, M. Subramanian, M. N. Islam, C. C. Rymond, J. Du, J. Hook, G. Kuriakose, J. Bhattacharya, I. Tabas, Regulatory T Cells Promote Macrophage Efferocytosis during Inflammation Resolution. Immunity (2018), doi:10.1016/j.immuni.2018.07.015.

13. C. B. Medina, P. Mehrotra, S. Arandjelovic, J. S. A. Perry, Y. Guo, S. Morioka, B. Barron, S. F. Walk, B. Ghesquière, A. S. Krupnick, U. Lorenz, K. S. Ravichandran, Metabolites released from apoptotic cells act as tissue messengers. Nature (2020), doi:10.1038/s41586-020-2121-3.

14. A. Trzeciak, Y. T. Wang, J. S. A. Perry, First we eat, then we do everything else: The dynamic metabolic regulation of efferocytosis. Cell Metab. 33, 2126–2141 (2021).

15. H. A. Anderson, C. A. Maylock, J. A. Williams, C. P. Paweletz, H. Shu, E. Shacter, Serum-derived protein S binds to phosphatidylserine and stimulates the phagocytosis of apoptotic cells. Nat. Immunol. 2002 41. 4, 87–91 (2002).

16. G. Lemke, Biology of the TAM Receptors. Cold Spring Harb. Perspect. Biol. 5, a009076 (2013).

17. Y. Belkaid, T. W. Hand, Role of the Microbiota in Immunity and inflammation. Cell. 157, 121 (2014).

18. E. Ansaldo, T. K. Farley, Y. Belkaid, Control of Immunity by the Microbiota. https://doi.org/10.1146/annurev-immunol-093019-112348. 39, 449–479 (2021).

19. Y. He, L. Fu, Y. Li, W. Wang, M. Gong, J. Zhang, X. Dong, J. Huang, Q. Wang, C. R. Mackay, Y. X. Fu, Y. Chen, X. Guo, Gut microbial metabolites facilitate anticancer therapy efficacy by modulating cytotoxic CD8+ T cell immunity. Cell Metab. 33, 988–1000.e7 (2021).

20. H. Liu, J. Wang, T. He, S. Becker, G. Zhang, D. Li, X. Ma, Butyrate: A Double-Edged Sword for Health? Adv. Nutr. 9, 21–29 (2018).

21. Y. Belkaid, S. Naik, Compartmentalized and systemic control of tissue immunity by commensals. Nat. Immunol. 2013 147. 14, 646–653 (2013).

22. G. Den Besten, K. Van Eunen, A. K. Groen, K. Venema, D. J. Reijngoud, B. M. Bakker, The role of short-chain fatty acids in the interplay between diet, gut microbiota, and host energy metabolism. J. Lipid Res. 54, 2325–2340 (2013).

23. J. H. Cummings, E. W. Pomare, H. W. J. Branch, C. P. E. Naylor, G. T. MacFarlane, Short chain fatty acids in human large intestine, portal, hepatic and venous blood. Gut. 28, 1221–1227 (1987).

24. J. Penders, I. Kummeling, C. Thijs, Infant antibiotic use and wheeze and asthma risk: a systematic review and meta-analysis. Eur. Respir. J. 38, 295–302 (2011).

25. L. Virta, A. Auvinen, H. Helenius, P. Huovinen, K. L. Kolho, Association of Repeated Exposure to Antibiotics With the Development of Pediatric Crohn’s Disease—A Nationwide, Register-based Finnish Case-Control Study. Am. J. Epidemiol. 175, 775–784 (2012).

26. R. W. Corty, B. W. Langworthy, J. P. Fine, J. B. Buse, H. K. Sanoff, J. L. Lund, Antibacterial Use Is Associated with an Increased Risk of Hematologic and Gastrointestinal Adverse Events in Patients Treated with Gemcitabine for Stage IV Pancreatic Cancer. Oncologist. 25, 579–584 (2020).

27. P. Nenclares, S. A. Bhide, H. Sandoval-Insausti, P. Pialat, L. Gunn, A. Melcher, K. Newbold, C. M. Nutting, K. J. Harrington, Impact of antibiotic use during curative treatment of locally advanced head and neck cancers with chemotherapy and radiotherapy. Eur. J. Cancer. 131, 9–15 (2020).

28. M. S. Martins Lopes, L. M. Machado, P. A. Ismael Amaral Silva, A. A. Tome Uchiyama, C. T. Yen, E. D. Ricardo, T. S. Mutao, J. R. Pimenta, D. S. Shimba, R. M. Hanriot, R. D. Peixoto, Antibiotics, cancer risk and oncologic treatment efficacy: a practical review of the literature. Ecancermedicalscience. 14, 1106 (2020).

29. S. Y. Shaw, J. F. Blanchard, C. N. Bernstein, Association Between the Use of Antibiotics and New Diagnoses of Crohnʼs Disease and Ulcerative Colitis. Am. J. Gastroenterol. 106, 2133–2142 (2011).

30. M. Norgaard, R. B. Nielsen, J. B. Jacobsen, J. L. Gradus, E. Stenager, N. Koch-Henriksen, T. L. Lash, H. T. Sorensen, Use of Penicillin and Other Antibiotics and Risk of Multiple Sclerosis: A Population-based Case-Control Study. Am. J. Epidemiol. 174, 945–948 (2011).

31. A. A. Sultan, C. Mallen, S. Muller, S. Hider, I. Scott, T. Helliwell, L. J. Hall, Antibiotic use and the risk of rheumatoid arthritis: a population-based case-control study. BMC Med. 17, 154 (2019).

32. L. H. Nguyen, A. K. Örtqvist, Y. Cao, T. G. Simon, B. Roelstraete, M. Song, A. D. Joshi, K. Staller, A. T. Chan, H. Khalili, O. Olén, J. F. Ludvigsson, Antibiotic use and the development of inflammatory bowel disease: a national case-control study in Sweden. Lancet Gastroenterol. Hepatol. (2020), doi:10.1016/s2468-1253(20)30267-3.

33. W. Solass, P. Horvath, F. Struller, I. Königsrainer, S. Beckert, A. Königsrainer, F.-J. Weinreich, M. Schenk, Functional vascular anatomy of the peritoneum in health and disease. Pleura and Peritoneum. 1, 145 (2016).

34. A. W. Roberts, B. L. Lee, J. Deguine, S. John, M. J. Shlomchik, G. M. Barton, Tissue-Resident Macrophages Are Locally Programmed for Silent Clearance of Apoptotic Cells. Immunity. 47, 913–927.e6 (2017).

35. S. Morioka, J. S. A. Perry, M. H. Raymond, C. B. Medina, Y. Zhu, L. Zhao, V. Serbulea, S. Onengut-Gumuscu, N. Leitinger, S. Kucenas, J. C. Rathmell, L. Makowski, K. S. Ravichandran, Efferocytosis induces a novel SLC program to promote glucose uptake and lactate release. Nature (2018),, doi:10.1038/s41586-018-0735-5.

36. P. Tirelle, J. Breton, G. Riou, P. Déchelotte, M. Coëffier, D. Ribet, Comparison of different modes of antibiotic delivery on gut microbiota depletion efficiency and body composition in mouse. BMC Microbiol. 20, 1–10 (2020).

37. A. Cifarelli, N. Forte, L. Lombardi, G. Pepe, F. Paradisi, The effect of some antibiotics on phagocytic activity in vitro. J. Infect. 5, 183–188 (1982).

38. S. Hodge, G. Hodge, S. Brozyna, H. Jersmann, M. Holmes, P. N. Reynolds, Azithromycin increases phagocytosis of apoptotic bronchial epithelial cells by alveolar macrophages. Eur. Respir. J. 28, 486–495 (2006).

39. R. Moges, D. D. De Lamache, S. Sajedy, B. S. Renaux, M. D. Hollenberg, G. Muench, E. M. Abbott, A. G. Buret, Anti-inflammatory benefits of antibiotics: Tylvalosin induces apoptosis of porcine neutrophils and macrophages, promotes efferocytosis, and inhibits pro-inflammatory CXCL-8, IL1α, and LTB4 production, while inducing the release of pro-resolving lipoxin A4 and resolvin D1. Front. Vet. Sci. 5, 57 (2018).

40. D. Han, M. C. Walsh, K. S. Kim, S. W. Hong, J. Lee, J. Yi, G. Rivas, C. D. Surh, Y. Choi, Microbiota-Independent Ameliorative Effects of Antibiotics on Spontaneous Th2-Associated Pathology of the Small Intestine. PLoS One. 10, e0118795 (2015).

41. H. G. Colaço, A. Barros, A. Neves-Costa, E. Seixas, D. Pedroso, T. Velho, K. L. Willmann, P. Faisca, G. Grabmann, H. S. Yi, M. Shong, V. Benes, S. Weis, T. Köcher, L. F. Moita, Tetracycline Antibiotics Induce Host-Dependent Disease Tolerance to Infection. Immunity. 54, 53–67.e7 (2021).

42. S. Gopinath, M. V. Kim, T. Rakib, P. W. Wong, M. Van Zandt, N. A. Barry, T. Kaisho, A. L. Goodman, A. Iwasaki, Topical application of aminoglycoside antibiotics enhances host resistance to viral infections in a microbiota-independent manner. Nat. Microbiol. 2018 35. 3, 611–621 (2018).

43. N. Moullan, L. Mouchiroud, X. Wang, D. Ryu, E. G. Williams, A. Mottis, V. Jovaisaite, M. V. Frochaux, P. M. Quiros, B. Deplancke, R. H. Houtkooper, J. Auwerx, Tetracyclines disturb mitochondrial function across eukaryotic models: a call for caution in biomedical research. Cell Rep. 10, 1681 (2015).

44. M. A. Woods Acevedo, J. K. Pfeiffer, Microbiota-Independent Antiviral Effects of Antibiotics on Poliovirus and Coxsackievirus. Virology. 546, 20 (2020).

45. E. A. Kennedy, K. Y. King, M. T. Baldridge, Mouse microbiota models: Comparing germ-free mice and antibiotics treatment as tools for modifying gut bacteria. Front. Physiol. 9, 1534 (2018).

46. Y. Okabe, R. Medzhitov, Tissue-specific signals control reversible program of localization and functional polarization of macrophages. Cell. 157, 832–844 (2014).

47. M. G. Constantinides, V. M. Link, S. Tamoutounour, A. C. Wong, P. J. Perez-Chaparro, S. J. Han, Y. E. Chen, K. Li, S. Farhat, A. Weckel, S. R. Krishnamurthy, I. Vujkovic-Cvijin, J. L. Linehan, N. Bouladoux, E. D. Merrill, S. Roy, D. J. Cua, E. J. Adams, A. Bhandoola, T. C. Scharschmidt, J. Aubé, M. A. Fischbach, Y. Belkaid, MAIT cells are imprinted by the microbiota in early life and promote tissue repair. Science (80-.). 366 (2019), doi:10.1126/SCIENCE.AAX6624/SUPPL_FILE/AAX6624-CONSTANTINIDES-SM.PDF.

48. A. Khosravi, A. Yáñez, J. G. Price, A. Chow, M. Merad, H. S. Goodridge, S. K. Mazmanian, Gut Microbiota Promote Hematopoiesis to Control Bacterial Infection. Cell Host Microbe. 15, 374–381 (2014).

49. G. A. Churchill, D. M. Gatti, S. C. Munger, K. L. Svenson, The Diversity Outbred Mouse Population. Mamm. Genome. 23, 713 (2012).

50. R. Oliveira Corrêa, J. L. Fachi, A. Vieira, F. T. Sato, M. Aurélio, R. Vinolo, Regulation of immune cell function by short-chain fatty acids. Clin. Transl. Immunol. 5, 73 (2016).

51. V. Sencio, A. Barthelemy, L. P. Tavares, M. G. Machado, D. Soulard, C. Cuinat, C. M. Queiroz-Junior, M. L. Noordine, S. Salomé-Desnoulez, L. Deryuter, B. Foligné, C. Wahl, B. Frisch, A. T. Vieira, C. Paget, G. Milligan, T. Ulven, I. Wolowczuk, C. Faveeuw, R. Le Goffic, M. Thomas, S. Ferreira, M. M. Teixeira, F. Trottein, Gut Dysbiosis during Influenza Contributes to Pulmonary Pneumococcal Superinfection through Altered Short-Chain Fatty Acid Production. Cell Rep. 30, 2934–2947.e6 (2020).

52. D. Erny, N. Dokalis, C. Mezö, A. Castoldi, O. Mossad, O. Staszewski, M. Frosch, M. Villa, V. Fuchs, A. Mayer, J. Neuber, J. Sosat, S. Tholen, O. Schilling, A. Vlachos, T. Blank, M. Gomez de Agüero, A. J. Macpherson, E. J. Pearce, M. Prinz, Microbiota-derived acetate enables the metabolic fitness of the brain innate immune system during health and disease. Cell Metab. 33, 2260–2276.e7 (2021).

53. M. H. Raymond, A. J. Davidson, Y. Shen, D. R. Tudor, C. D. Lucas, S. Morioka, J. S. A. Perry, J. Krapivkina, D. Perrais, L. J. Schumacher, R. E. Campbell, W. Wood, K. S. Ravichandran, Live cell tracking of macrophage efferocytosis during Drosophila embryo development in vivo. Science (80-.). 375, 1182–1187 (2022).

54. A. J. Davidson, W. Wood, Macrophages Use Distinct Actin Regulators to Switch Engulfment Strategies and Ensure Phagocytic Plasticity In Vivo. Cell Rep. 31, 107692 (2020).

55. S. Uderhardt, A. J. Martins, J. S. Tsang, T. Lämmermann, R. N. Germain, Resident Macrophages Cloak Tissue Microlesions to Prevent Neutrophil-Driven Inflammatory Damage. Cell. 177, 541–555.e17 (2019).

56. J. Schulthess, S. Pandey, M. Capitani, K. C. Rue-Albrecht, I. Arnold, F. Franchini, A. Chomka, N. E. Ilott, D. G. W. Johnston, E. Pires, J. McCullagh, S. N. Sansom, C. V. Arancibia-Cárcamo, H. H. Uhlig, F. Powrie, The Short Chain Fatty Acid Butyrate Imprints an Antimicrobial Program in Macrophages. Immunity (2019), doi:10.1016/j.immuni.2018.12.018.

57. M. R. Elliott, K. S. Ravichandran, The Dynamics of Apoptotic Cell Clearance. Dev. Cell (2016),, doi:10.1016/j.devcel.2016.06.029.

58. C. Lantz, B. Radmanesh, E. Liu, E. B. Thorp, J. Lin, Single-cell RNA sequencing uncovers heterogenous transcriptional signatures in macrophages during efferocytosis. Sci. Reports 2020 101. 10, 1–11 (2020).

59. T. S. P. Heng, M. W. Painter, K. Elpek, V. Lukacs-Kornek, N. Mauermann, S. J. Turley, D. Koller, F. S. Kim, A. J. Wagers, N. Asinovski, S. Davis, M. Fassett, M. Feuerer, D. H. D. Gray, S. Haxhinasto, J. A. Hill, G. Hyatt, C. Laplace, K. Leatherbee, D. Mathis, C. Benoist, R. Jianu, D. H. Laidlaw, J. A. Best, J. Knell, A. W. Goldrath, J. Jarjoura, J. C. Sun, Y. Zhu, L. L. Lanier, A. Ergun, Z. Li, J. J. Collins, S. A. Shinton, R. R. Hardy, R. Friedline, K. Sylvia, J. Kang, The Immunological Genome Project: networks of gene expression in immune cells. Nat. Immunol. 2008 910. 9, 1091–1094 (2008).

60. C. González-Bosch, E. Boorman, P. A. Zunszain, G. E. Mann, Short-chain fatty acids as modulators of redox signaling in health and disease. Redox Biol. 47, 102165 (2021).

61. B. van der Hee, J. M. Wells, Microbial Regulation of Host Physiology by Short-chain Fatty Acids. Trends Microbiol. 29, 700–712 (2021).

62. R. Corrêa-Oliveira, J. L. Fachi, A. Vieira, F. T. Sato, M. A. R. Vinolo, Regulation of immune cell function by short-chain fatty acids. Clin. Transl. Immunol. 5, e73 (2016).

63. C. Wu, I. MacLeod, A. I. Su, BioGPS and MyGene.info: organizing online, gene-centric information. Nucleic Acids Res. 41, D561–D565 (2013).

64. A. Nakajima, A. Nakatani, S. Hasegawa, J. Irie, K. Ozawa, G. Tsujimoto, T. Suganami, H. Itoh, I. Kimura, The short chain fatty acid receptor GPR43 regulates inflammatory signals in adipose tissue M2-type macrophages. PLoS One. 12, e0179696 (2017).

65. R. Schilderink, C. Verseijden, J. Seppen, V. Muncan, G. R. van den Brink, T. T. Lambers, E. van Tol, W. J. de Jonge, The SCFA butyrate stimulates the epithelial production of retinoic acid via inhibition of epithelial HDAC. Am. J. Physiol. - Gastrointest. Liver Physiol. 310, G1138–G1146 (2016).

66. S. Löfmark, C. Jernberg, J. K. Jansson, C. Edlund, Clindamycin-induced enrichment and long-term persistence of resistant Bacteroides spp. and resistance genes. J. Antimicrob. Chemother. 58, 1160–1167 (2006).

67. C. Jernberg, S. Löfmark, C. Edlund, J. K. Jansson, Long-term ecological impacts of antibiotic administration on the human intestinal microbiota. ISME J. 2007 11. 1, 56–66 (2007).

68. H. E. Jakobsson, C. Jernberg, A. F. Andersson, M. Sjölund-Karlsson, J. K. Jansson, L. Engstrand, Short-Term Antibiotic Treatment Has Differing Long-Term Impacts on the Human Throat and Gut Microbiome. PLoS One. 5, e9836 (2010).

69. K. Korpela, A. Salonen, L. J. Virta, R. A. Kekkonen, K. Forslund, P. Bork, W. M. De Vos, Intestinal microbiome is related to lifetime antibiotic use in Finnish pre-school children. Nat. Commun. 2016 71. 7, 1–8 (2016).

70. L. Dethlefsen, S. Huse, M. L. Sogin, D. A. Relman, The Pervasive Effects of an Antibiotic on the Human Gut Microbiota, as Revealed by Deep 16S rRNA Sequencing. PLOS Biol. 6, e280 (2008).

71. R. Vangoitsenhoven, G. A. M. Cresci, Role of Microbiome and Antibiotics in Autoimmune Diseases. Nutr. Clin. Pract. 35, 406–416 (2020).

72. C. V. Rothlin, T. D. Hille, S. Ghosh, Determining the effector response to cell death. Nat. Rev. Immunol. 2020 215. 21, 292–304 (2020).

73. J. Newson, M. Stables, E. Karra, F. Arce-Vargas, S. Quezada, M. Motwani, M. Mack, S. Yona, T. Audzevich, D. W. Gilroy, Resolution of acute inflammation bridges the gap between innate and adaptive immunity. Blood. 124, 1748–1764 (2014).

74. I. Cho, S. Yamanishi, L. Cox, B. A. Methé, J. Zavadil, K. Li, Z. Gao, D. Mahana, K. Raju, I. Teitler, H. Li, A. V. Alekseyenko, M. J. Blaser, Antibiotics in early life alter the murine colonic microbiome and adiposity. Nat. 2012 4887413. 488, 621–626 (2012).

75. J. Henao-Mejia, E. Elinav, C. Jin, L. Hao, W. Z. Mehal, T. Strowig, C. A. Thaiss, A. L. Kau, S. C. Eisenbarth, M. J. Jurczak, J. P. Camporez, G. I. Shulman, J. I. Gordon, H. M. Hoffman, R. A. Flavell, Inflammasome-mediated dysbiosis regulates progression of NAFLD and obesity. Nat. 2012 4827384. 482, 179–185 (2012).

76. C. Canova, V. Zabeo, G. Pitter, P. Romor, T. Baldovin, R. Zanotti, L. Simonato, Association of Maternal Education, Early Infections, and Antibiotic Use With Celiac Disease: A Population-Based Birth Cohort Study in Northeastern Italy. Am. J. Epidemiol. 180, 76–85 (2014).

77. L. M. Cox, S. Yamanishi, J. Sohn, A. V. Alekseyenko, J. M. Leung, I. Cho, S. G. Kim, H. Li, Z. Gao, D. Mahana, J. G. Zárate Rodriguez, A. B. Rogers, N. Robine, P. Loke, M. J. Blaser, Altering the Intestinal Microbiota during a Critical Developmental Window Has Lasting Metabolic Consequences. Cell. 158, 705–721 (2014).

78. L. M. Cox, M. J. Blaser, Pathways in Microbe-Induced Obesity. Cell Metab. 17, 883–894 (2013).

79. K. M. Kemppainen, K. Vehik, K. F. Lynch, H. E. Larsson, R. J. Canepa, V. Simell, S. Koletzko, E. Liu, O. G. Simell, J. Toppari, A. G. Ziegler, M. J. Rewers, Å. Lernmark, W. A. Hagopian, J. X. She, B. Akolkar, D. A. Schatz, M. A. Atkinson, M. J. Blaser, J. P. Krischer, H. Hyöty, D. Agardh, E. W. Triplett, K. Bautista, J. Baxter, R. Bedoy, D. Felipe-Morales, K. Driscoll, B. I. Frohnert, P. Gesualdo, M. Hoffman, R. Karban, J. Norris, A. Samper-Imaz, A. Steck, K. Waugh, H. Wright, A. Adamsson, S. Ahonen, J. Ilonen, S. Jokipuu, T. Kallio, L. Karlsson, M. Kähönen, M. Knip, L. Kovanen, M. Koreasalo, K. Kurppa, T. Latva-Aho, M. Lönnrot, E. Mäntymäki, K. Multasuo, J. Mykkänen, T. Niininen, S. Niinistö, M. Nyblom, P. Rajala, J. Rautanen, A. Riikonen, M. Riikonen, J. Rouhiainen, M. Romo, T. Simell, M. Sjöberg, A. Stenius, M. Leppänen, S. Vainionpää, E. Varjonen, R. Veijola, S. M. Virtanen, M. Vähä-Mäkilä, M. Åkerlund, K. Lindfors, D. Hopkins, L. Steed, J. Thomas, J. Adams, K. Silvis, M. Haller, M. Gardiner, R. McIndoe, A. Sharma, J. William, G. Young, S. W. Anderson, L. Jacobsen, A. Beyerlein, E. Bonifacio, M. Hummel, S. Hummel, K. Foterek, N. Janz, M. Kersting, A. Knopff, C. Peplow, R. Roth, M. Scholz, J. Stock, K. Warncke, L. Wendel, C. Winkler, C. A. Aronsson, M. Ask, J. Bremer, U. M. Carlsson, C. Cilio, E. Ericson-Hallström, L. Fransson, T. Gard, J. Gerardsson, R. Bennet, M. Hansen, G. Hansson, S. Hyberg, F. Johansen, B. Jonsdottir, M. Lindström, M. Lundgren, M. Månsson-Martinez, M. Markan, J. Melin, Z. Mestan, K. Ottosson, K. Rahmati, A. Ramelius, F. Salami, S. Sibthorpe, B. Sjöberg, U. Swartling, E. T. Amboh, C. Törn, A. Wallin, Å. Wimar, S. Åberg, M. Killian, C. C. Crouch, J. Skidmore, J. Carson, M. Dalzell, K. Dunson, R. Hervey, C. Johnson, R. Lyons, A. Meyer, D. Mulenga, A. Tarr, M. Uland, J. Willis, D. Becker, M. Franciscus, M. E. D. E. Smith, A. Daftary, M. B. Klein, C. Yates, M. Abbondondolo, S. Austin-Gonzalez, M. Avendano, S. Baethke, R. Brown, B. Burkhardt, M. Butterworth, J. Clasen, D. Cuthbertson, C. Eberhard, S. Fiske, D. Garcia, J. Garmeson, V. Gowda, K. Heyman, F. P. Laras, H. S. Lee, S. Liu, X. Liu, K. Lynch, J. Malloy, C. McCarthy, S. Meulemans, H. Parikh, C. Shaffer, L. Smith, S. Smith, N. Sulman, R. Tamura, U. Uusitalo, P. Vijayakandipan, K. Wood, J. Yang, L. Ballard, D. Hadley, W. McLeod, K. Bourcier, T. Briese, S. B. Johnson, L. Yu, D. Miao, P. Bingley, A. William’s, K. Chandler, S. Rokni, C. Willia, R. Wyatt, G. George, S. Grace, N. Mulholland, Association Between Early-Life Antibiotic Use and the Risk of Islet or Celiac Disease Autoimmunity. JAMA Pediatr. 171, 1217–1225 (2017).

80. P. V. Chang, L. Hao, S. Offermanns, R. Medzhitov, The microbial metabolite butyrate regulates intestinal macrophage function via histone deacetylase inhibition. Proc. Natl. Acad. Sci. U. S. A. 111, 2247–2252 (2014).

81. M. R. Fernando, A. Saxena, J. L. Reyes, D. M. McKay, Butyrate enhances antibacterial effects while suppressing other features of alternative activation in IL-4-induced macrophages. Am. J. Physiol. - Gastrointest. Liver Physiol. 310, G822–G831 (2016).

82. C. C. Bain, A. Bravo-Blas, C. L. Scott, E. Gomez Perdiguero, F. Geissmann, S. Henri, B. Malissen, L. C. Osborne, D. Artis, A. M. I. Mowat, Constant replenishment from circulating monocytes maintains the macrophage pool in the intestine of adult mice. Nat. Immunol. 2014 1510. 15, 929–937 (2014).

83. J. A. Nicolás-Ávila, A. V. Lechuga-Vieco, L. Esteban-Martínez, M. Sánchez-Díaz, E. Díaz-García, D. J. Santiago, A. Rubio-Ponce, J. L. Y. Li, A. Balachander, J. A. Quintana, R. Martínez-de-Mena, B. Castejón-Vega, A. Pun-García, P. G. Través, E. Bonzón-Kulichenko, F. García-Marqués, L. Cussó, N. A-González, A. González-Guerra, M. Roche-Molina, S. Martin-Salamanca, G. Crainiciuc, G. Guzmán, J. Larrazabal, E. Herrero-Galán, J. Alegre-Cebollada, G. Lemke, C. V. Rothlin, L. J. Jimenez-Borreguero, G. Reyes, A. Castrillo, M. Desco, P. Muñoz-Cánoves, B. Ibáñez, M. Torres, L. G. Ng, S. G. Priori, H. Bueno, J. Vázquez, M. D. Cordero, J. A. Bernal, J. A. Enríquez, A. Hidalgo, A Network of Macrophages Supports Mitochondrial Homeostasis in the Heart. Cell. 183, 94–109.e23 (2020).

84. J. R. Wright, Clearance and recycling of pulmonary surfactant. https://doi.org/10.1152/ajplung.1990.259.2.L1. 259 (1990), doi:10.1152/AJPLUNG.1990.259.2.L1.

85. K. K. Penberthy, J. J. Lysiak, K. S. Ravichandran, Rethinking Phagocytes: Clues from the Retina and Testes. Trends Cell Biol. 28, 317–327 (2018).

86. J. S. A. Perry, S. Morioka, C. B. Medina, J. Iker Etchegaray, B. Barron, M. H. Raymond, C. D. Lucas, S. Onengut-Gumuscu, E. Delpire, K. S. Ravichandran, Interpreting an apoptotic corpse as anti-inflammatory involves a chloride sensing pathway. Nat. Cell Biol. (2019), doi:10.1038/s41556-019-0431-1.

