## Supplementary Figures for "A microbiota-derived metabolite instructs peripheral efferocytosis"

### Supplementary Figures 1 – 6

**Fig. S1. The intestinal microbiome supports peripheral efferocytosis.**

(A) Representative flow cytometry plots of $CD11b^+F4/80^+$ cells from peritoneal lavage of mice following antibiotics treatment for indicated days (Day 0 = blue, Day 3 = light green, Day 7 = dark green).

(B) Percentage of $CD11b^+F4/80^+$ macrophages isolated from peritoneal lavage in (A). *$p < .05$.

(C) Percentage of $CD11b^+F4/80^+$ peritoneal macrophages from SPF (blue) and GF (purple) mice. ns = not significant.

(D) *In vitro* efferocytosis analysis of BMDMs from SPF or GF mice. ns = not significant.

(E) Representative flow cytometry plots of PI and AnnexinV staining of thymocytes 6 hours after dexamethasone injection.

(F) Percentage of $CD11b^+F4/80^+$ peritoneal macrophages from C57BL6/J (blue) and J:DO (brown) mice. ****$p < .0001$.

### Fig. S1: The intestinal microbiome supports peripheral efferocytosis

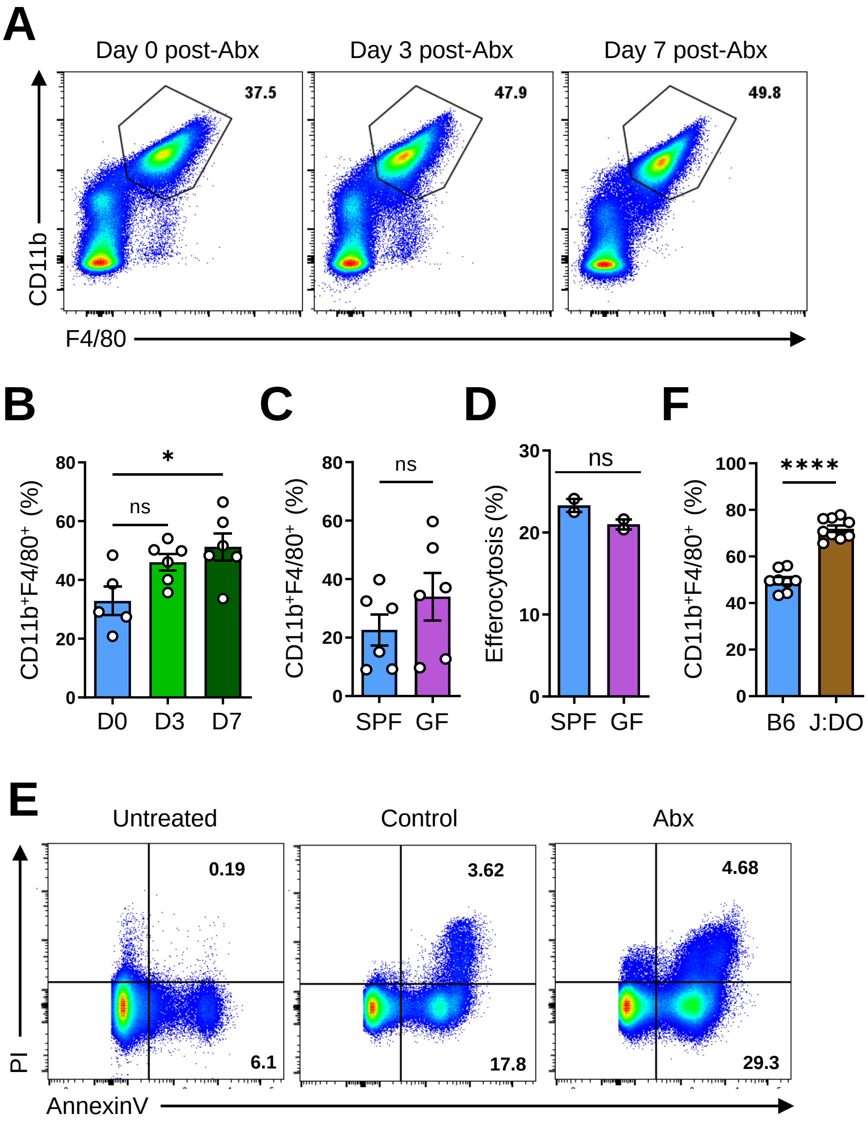

**Fig. S2. Butyrate boosts efferocytosis via induction of efferocytotic transcriptional programs.**

(A) Experimental protocol for experiments performed in **Fig. 2C** and **Fig. 2D**.

(B) Summary data from flow cytometry analysis of *in vitro* efferocytosis by BMDMs treated with vehicle (white) or butyrate (orange, 1 mM) for 16h. ns = not significant.

(C) Representative flow cytometry plots of vehicle and butyrate (1 mM)-conditioned BMDMs for the indicated time points from experiments shown in **Fig. 2D**. In some conditions, BMDMs were conditioned for 3d followed by withdraw of butyrate for the indicated time point prior to efferocytosis assay.

(D) Representative transluminescence microscopy image of vehicle and butyrate-treated BMDMs from experiments performed in **Fig. 2C**.

(E) Flow cytometry analysis and quantification of *in vitro* antibody-mediated phagocytosis in BMDMs conditioned with vehicle or butyrate (1 mM) for 3d. ****$p$ < .0001.

(F) Efferocytosis-associated programs identified in butyrate-treated BMDMs from experiments performed as in **Fig. 2G**. Data corresponds to analyses presented in **Fig. 2H**.

(G) Putative efferocytosis-supporting programs identified in butyrate-treated BMDMs from experiments performed as in **Fig. 2G**. Data corresponds to analyses presented in **Fig. 2I**.

**Fig. S2: Butyrate boosts efferocytosis via induction of efferocytotic transcriptional programs**

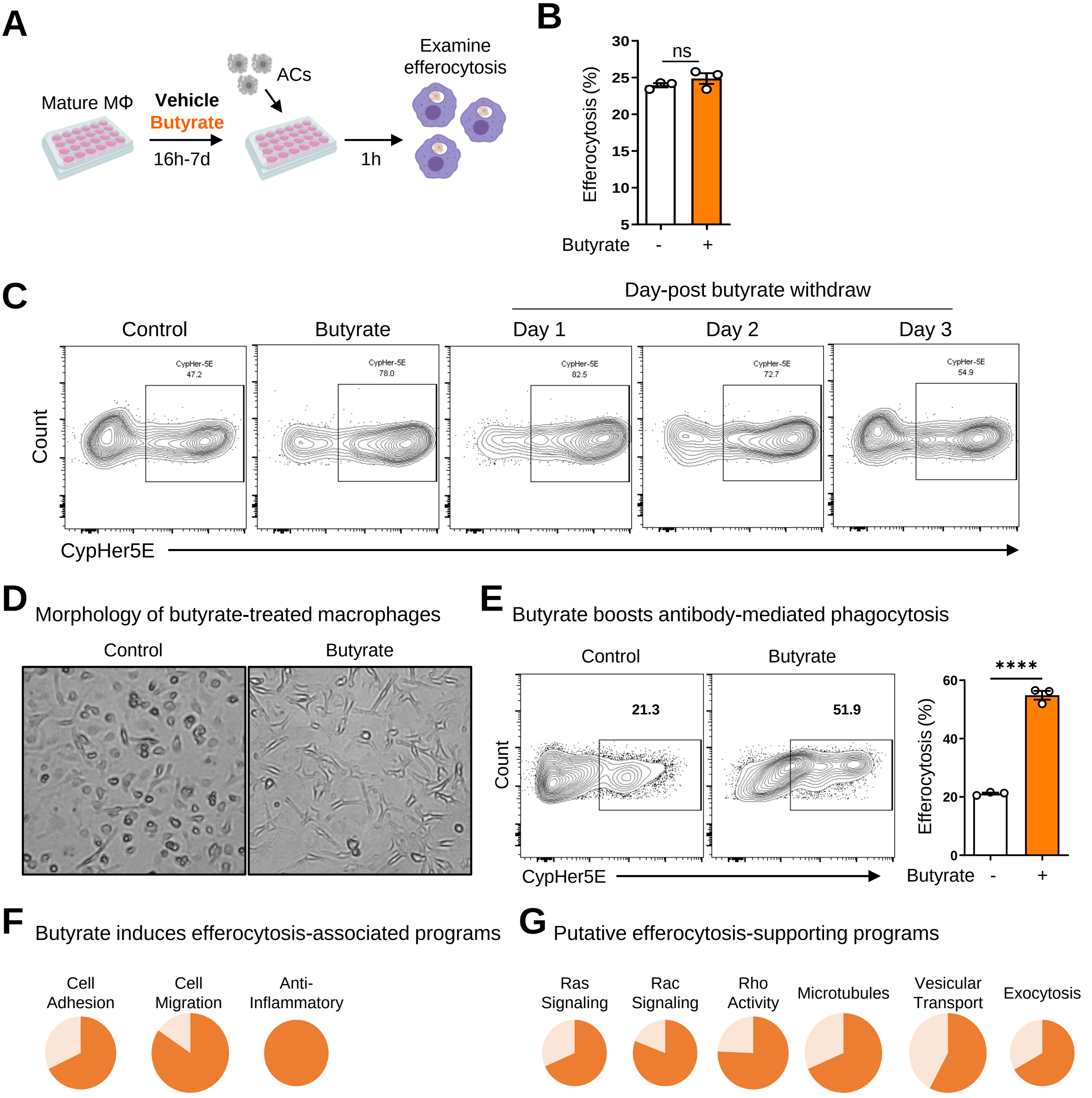

**Fig. S3. Exogenous butyrate rescues antibiotic-induced defects in peripheral efferocytosis.** Shown are the UMAP representations of total peritoneal cells and top uniquely differentially expressed genes for each cluster from scRNAseq experiments performed as in **Fig. 3C**.

### Fig. S3: Exogenous butyrate rescues antibiotic-induced defects in peripheral efferocytosis

UMAP and top uniquely upregulated genes from scRNAseq of peritoneal cells in mice treated as indicated

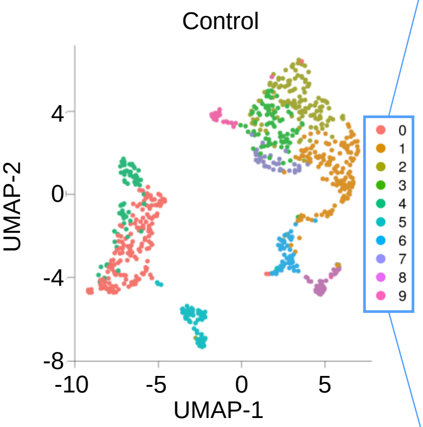


| Cluster 0 | Cluster 1 | Cluster 2 | Cluster 3 | Cluster 4 |
|---|---|---|---|---|
| *Ms4a1* | *Fn1* | *Hp* | *Selenof* | *Ighd* |
| *Igcl2* | *Itgam* | *Sntb2* | *Atp6v0e* | *Ccr7* |
| *Ighm* | *Gda* | *Pi16* | *Tspan4* | |
| *Cd79b* | *Sdc3* | *Ctla2b* | *Lamtor3* | |
| *Ly6d* | *Emilin2* | *Isyna1* | *Psmb2* | |

| Cluster 5 | Cluster 6 | Cluster 8 |
|---|---|---|
| *Ms4a4b* | *Ltb4r1* | *Cdk1* |
| *Trbc2* | *Clec4b1* | *Birc5* |
| *Il7r* | *Ccr2* | *Nusap1* |
| *Cd3d* | *Trerf1* | *Cenpf* |
| *Cd3g* | *Plbd1* | *Cdca3* |

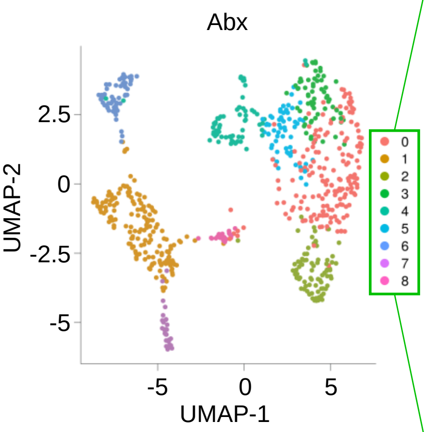


| Cluster 0 | Cluster 1 | Cluster 2 | Cluster 3 | Cluster 4 |
|---|---|---|---|---|
| *Prg4* | *Ebf1* | *Marco* | *Cd81* | *Spc24* |
| *Selp* | *Cd79a* | *Slc7a2* | *Bin1* | *Tk1* |
| *Fam46a* | *Ly6d* | *Grn* | *Pfkl* | *Cdk1* |
| *Fn1* | *Ms4a1* | *Ahnak* | *Pla2g7* | *Mki67* |
| *Itgam* | *Igcl2* | *Vps13c* | *Fcgr1* | *Stmn1* |

| Cluster 5 | Cluster 6 | Cluster 8 |
|---|---|---|
| *Fth1* | *Il7r* | *Clec4b1* |
| | *Ms4a4b* | *Plbd1* |
| | *Trbc2* | *Ltb4r1* |
| | *Cd3d* | *Cd209a* |
| | *Txk* | *Ccr2* |

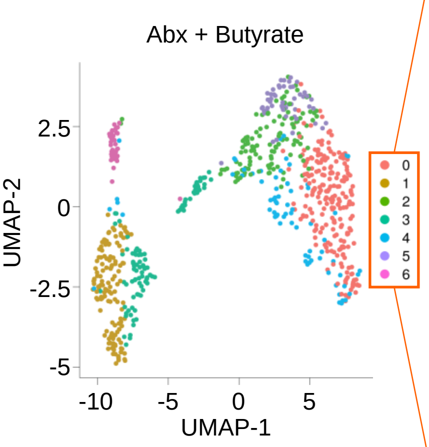


| Cluster 0 | Cluster 1 | Cluster 2 | Cluster 3 |
|---|---|---|---|
| *Prg4* | *Plac8* | *Chil3* | *Mki67* |
| *Fn1* | *Ms4a1* | *Cd81* | *Stmn1* |
| *Cfh* | *Mzb1* | *Chchd2* | *Top2a* |
| *F5* | *Ly6a* | *Ubl5* | *Lig1* |
| *Itgam* | *Ly6d* | *Atp5h* | *Smc2* |

| Cluster 4 | Cluster 6 |
|---|---|
| *Aif1* | *Il7r* |
| *Ptpn12* | *Cd3d* |
| *H2-dmb1* | *Trbc2* |
| *Gm2a* | *Ms4a4b* |
| *Wars* | *Trbc1* |

**Fig. S4. Exogenous butyrate rescues antibiotic-induced defects in peripheral efferocytosis.** Similar to Fig. S3, except showing overlays of key cell type identifiers for immune cell subtypes in scRNAseq data from **Fig. 3C**. *Itgam* and *Fcer1g* are found in large peritoneal macrophages (LPMs), *Clec4b1* and *Ltb4r1* are found in small peritoneal macrophages (SPMs), *Cd79b* and *Ms4a1* are found in B1 B cells, and *Trbc2* and *Cd3d* are found in T cells.

**Fig. S4: Exogenous butyrate rescues antibiotic-induced defects in peripheral efferocytosis**

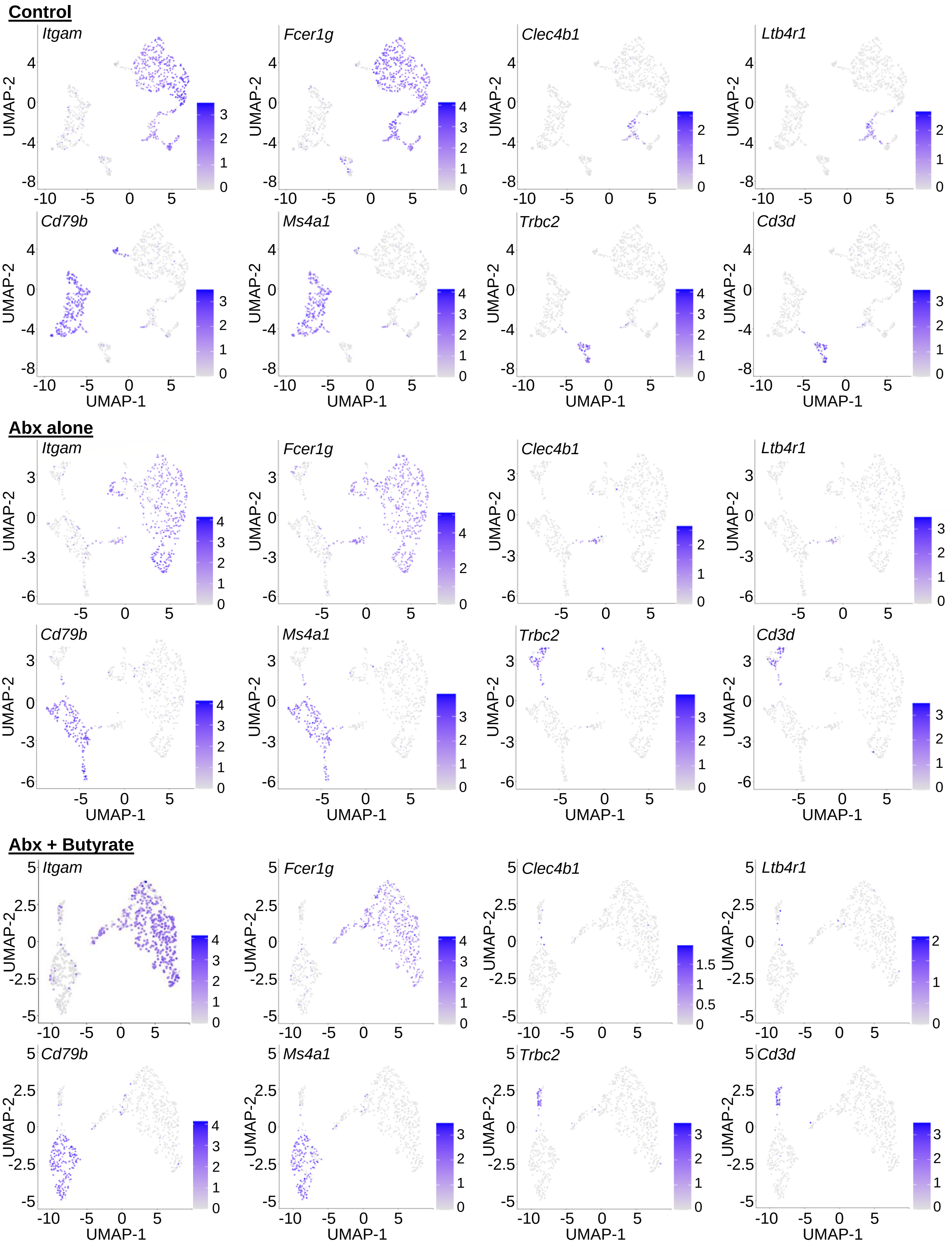

**Fig. S5. Butyrate boosts efferocytosis via histone acetylation, not cognate GPCR signaling.**

(A, B) Percentage of $CD11b^+F4/80^+$ peritoneal macrophages from *Ffar2*$^{-/-}$ (A) (n = 3) and *Hcar2*$^{-/-}$ (B) (n = 5) mice. ns = not significant.

(C) Efferocytosis by peritoneal macrophages does not depend on GPR43 *in vivo*. *Ffar2*$^{-/-}$ (n=5) mice were injected intraperitoneally (i.p.) with CypHer5E-labeled ACs. After 1h, peritoneal $CD11b^+F4/80^+$ macrophages were analyzed for efferocytosis. Shown are representative flow cytometry (left) and summary (right) plots of the rate of efferocytosis by $CD11b^+F4/80^+$ macrophages. ns = not significant. Data are from two independent experiments.

(D) Representative flow cytometry plots of acetylation and methylation of the indicated histone 3 lysine in BMDMs treated with vehicle (black) or butyrate for 3d (orange, 1 mM). Values shown within histograms represent median fluorescence intensity.

#### Fig. S5: Butyrate boosts efferocytosis via histone acetylation, not cognate GPCR signaling

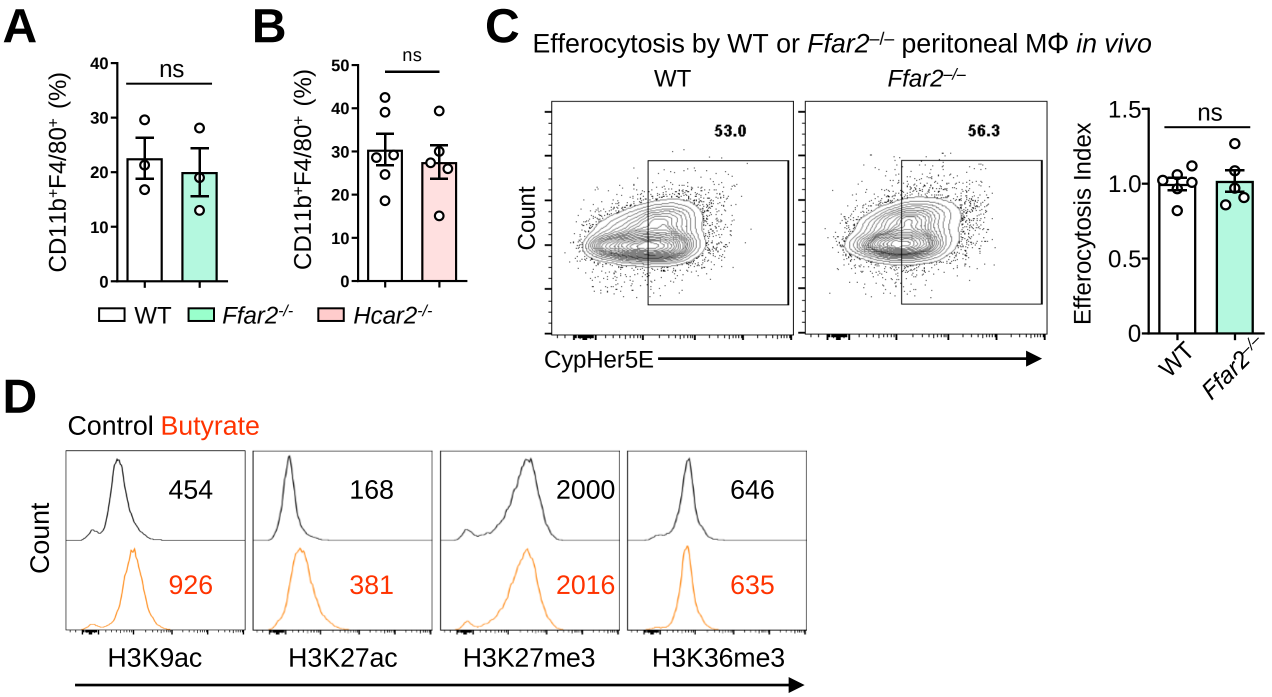

**Fig. S6. Treatment with antibiotics induces prolonged peripheral efferocytosis defects.** Representative flow cytometry plots of F4/80 (macrophage) and Ly6G (neutrophil) staining of peritoneal cells 3d post-injection of zymosan.

**Fig. S6: Treatment with antibiotics induces prolonged peripheral efferocytosis defects**

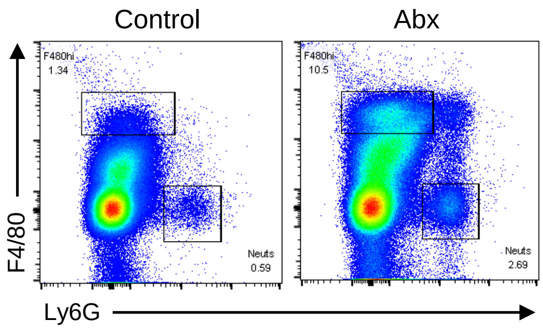
